# Genomic trinucleotide composition reflects context-dependent mutation spectra in plants and mammals

**DOI:** 10.64898/2026.11.23.768609

**Authors:** Yijie Wang, Ziyue Gao

**Affiliations:** Department of Genetics, Perelman School of Medicine, University of Pennsylvania, Philadelphia, PA 19104, USA

## Abstract

Mutation rates vary substantially across mutation types and sequence contexts, yet it remains unclear how variation in the mutation spectrum shapes genome composition across species. Motivated by the strong cross-species relationship between CpG mutation rate and CpG content, we hypothesize that multi-nucleotide genome composition reflects the context-dependent mutation spectrum, with additional effects of GC-biased gene conversion (gBGC). To test this hypothesis, we develop a framework to determine the equilibrium trinucleotide composition from a context-dependent mutation spectrum, with and without incorporating the effects of gBGC. The trinucleotide frequencies predicted by our model based on polymorphism-derived mutation spectra of 25 plants and 10 mammals show good concordance with observed genomic trinucleotide compositions, and incorporating gBGC further improves agreement across all species, highlighting the pervasive influence of gBGC on genome composition. We further find that genome composition retains information about lineage-specific context-dependent mutation spectra, with strong correspondence within evolutionary clades but limited species-level specificity. In particular, trinucleotide abundance ratios correlate positively with the corresponding reciprocal mutation rate ratios, both across trinucleotide pairs within species and across species for the same trinucleotide pair. Together, our results demonstrate that genome composition encodes quantitative information about mutation processes and suggest that a single reference genome can inform mutational patterns in species lacking polymorphism data or direct measurements of mutation.

## Introduction

Genome composition—the frequencies of individual nucleotides or short sequence motifs—is one of the most conspicuous and variable features of genomes. GC content varies markedly among species, chromosomes, and genomic regions, and dinucleotide and other oligonucleotide frequencies provide additional compositional signatures that can distinguish species even after differences in overall base composition are considered (1-3). A prominent example is the depletion of CpG dinucleotides in many methylated genomes, which reflects the elevated rate at which methylated cytosines mutate to thymine (4). Composition also differs among coding, regulatory, repetitive, and intergenic sequences because these genomic compartments experience different molecular processes as well as functional and evolutionary constraints (2, 3). Genome composition is therefore not merely a descriptive property: it is an evolutionary record of mutational input and the forces that govern the fate of new genetic variants.

Mutation provides the initial input into this process. Single-nucleotide mutation rates differ among substitution types and depend strongly on the surrounding sequence (5-7). Mutation spectra summarize these biases by recording the relative rates of different nucleotide changes in their local contexts. A widely used representation folds strand-complementary changes into 96 categories, corresponding to six possible substitutions of the central base and 16 combinations of its immediate 5′ and 3′ flanking nucleotides (8, 9). Although this trinucleotide representation does not capture all effects of extended sequence context (7, 10-13), it captures major mutational processes, including the high mutability of methylated CpG sites (4, 11). Importantly, mutation spectra vary among populations and species, potentially reflecting distinct DNA modification, replication, damage, and repair processes or differential activities of shared processes (5, 9, 11, 14-17).

The long-term effects of mutation on genome composition can be formalized using mutation equilibrium theory. Under a stable mutation process, sequence composition is expected to approach an equilibrium at which mutational flux into each sequence state balances flux out of it (18, 19). In the simplest nucleotide model, equilibrium GC content is determined by the relative rates of mutations from A/T to G/C and from G/C to A/T. Extending this principle to trinucleotides is more complex because neighboring bases influence mutation rates, while each mutation alters the sequence contexts of multiple overlapping trinucleotides. Equilibrium trinucleotide composition needs to therefore be modeled as a coupled system of context-dependent transitions rather than by considering each mutation type independently.

Mutation equilibrium provides a useful baseline model, but mutation alone does not determine genome composition. Once a mutation arises, its contribution to long-term genome composition depends on the probability that it is transmitted and ultimately fixed. One important influence on this process is GC-biased gene conversion (gBGC), a recombination-associated transmission bias that preferentially favors G/C alleles over A/T alleles during the repair of mismatches in meiotic heteroduplex DNA (20). Although mechanistically distinct from natural selection, gBGC alters the fixation probabilities of mutations between A/T and G/C in a manner analogous to natural selection (20, 21). Consistent with this effect, observed genomic GC content often exceeds the equilibrium predicted from mutation alone (22-24). Moreover, regional changes in GC content are frequently correlated with variation in recombination rate (20, 25, 26). Comparative studies further suggest that gBGC has shaped genome composition across diverse animals and plants (27-29).

Despite extensive work on these processes that shape genome composition, their relative contributions to and interactive effects on multi-nucleotide composition remain poorly understood. Most previous studies of genome composition evolution have focused on overall GC content, regional GC heterogeneity, or dinucleotide relative abundance (1-3, 20). Conversely, studies of context-dependent mutation have generally described mutation spectra without asking whether those spectra quantitatively generate the observed frequencies of short sequence motifs (5, 11, 15, 17). It therefore remains unclear how much variation in trinucleotide composition can be explained by mutation alone and how much reflects gBGC or other evolutionary forces. A related question is whether this relationship can be read in reverse: although genome composition cannot capture every feature of the underlying mutation spectrum, it may retain broad signatures of lineage-specific mutational processes (11). Addressing these questions requires a framework that connects context-dependent mutation rates to equilibrium multi-nucleotide composition while allowing the effects of mutation and fixation biases to be evaluated separately.

Here, we characterize intergenic trinucleotide composition across 88 plant and mammalian genomes and develop an iterative discrete-time Markov-chain framework for predicting stationary trinucleotide frequencies from context-dependent mutation spectra. For 35 species with polymorphism-based mutation spectra, we compare observed composition with predictions from mutation alone and from a model that incorporates gBGC as a fixation bias. We further test whether trinucleotide composition reflects differences in mutation spectra across species. We find that contemporary mutation spectra explain much of the observed trinucleotide composition, that incorporating gBGC substantially improves agreement for most species, and that genome composition retains quantitative, lower-dimensional information about lineage-specific mutational processes. These results establish a mechanistic link between context-dependent mutation, fixation bias, and the long-term evolution of genome composition.

## Results

### Cross-species variation in trinucleotide composition is dominated by GC content and CpG depletion

To characterize the shared and distinct features of genomic multi-nucleotide composition across species, we analyzed the trinucleotide (3-mer) composition across 75 plant and 13 mammalian genomes (Dataset S1), after removing low complexity regions and simple repeats to reduce technical issues, and genic regions to minimize effect of direct purifying selection (Fig. 1A; Figs S1-S3; Dataset S2; Methods). Because transposable elements (TEs) comprise highly variable fractions of plant genomes, we also evaluated multiple TE-masking strategies in representative species (Methods). Although TE masking removed substantial amounts of sequence, the resulting trinucleotide compositions were highly similar to those obtained without TE masking (Jensen-Shannon divergence ≤ 0.003; Fig. S4) with somewhat greater differences in maize, where more than 80% of genomic sequences are TEs (Fig. S4C). We therefore retained TEs in the primary analyses to maximize the amount of sequence analyzed.

**Figure 1.**
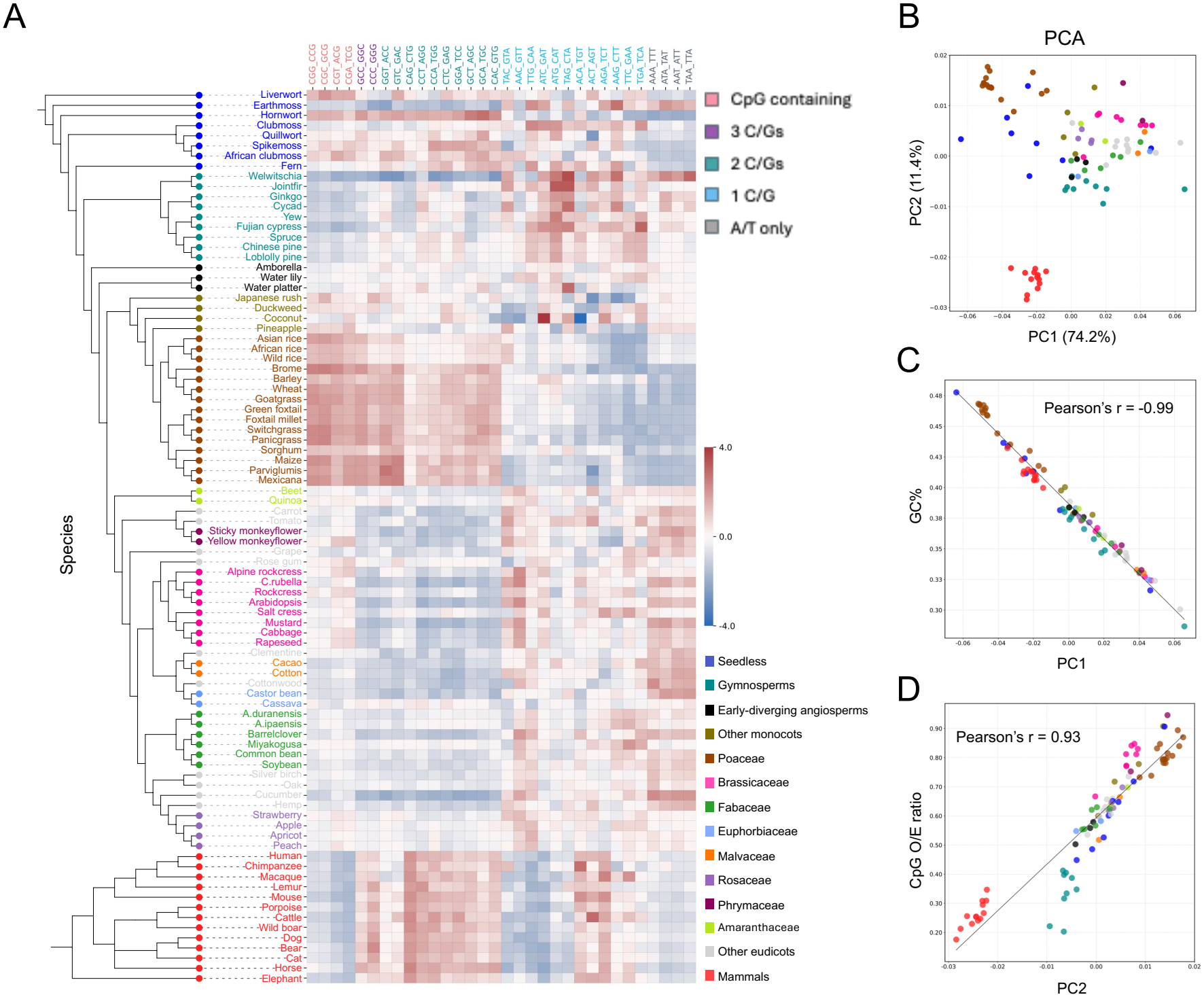
Variation in trinucleotide genome composition across 75 plant and 13 mammal species is primarily characterized by GC content and CpG depletion. **A**. Heatmap of normalized trinucleotide frequencies in intergenic regions after collapsing reverse-complement pairs (e.g., ACC and GGT). Colors of species names indicate broad taxonomic groups; colors of trinucleotides indicate GC content and CpG-containing status. Color coding for taxonomic groups and trinucleotide classes is consistent throughout all figures. Z-score normalization was performed across species separately for each trinucleotide, such that heatmap colors represent the relative abundance of a trinucleotide across species. **B**. PCA of trinucleotide frequencies in intergenic regions across 88 species. In **B–D**, each point represents one species, with colors indicating the same broad taxonomic groups as in A. **C–D**. Correlations between GC% and PC1 (**C**) and between CpG observed-to-expected (O/E) ratio and PC2 (**D**).

Several prominent features of trinucleotide composition were shared broadly across species. CpG-containing 3-mers tend to be among the least abundant 3-mers (Fig. S5) and consistently depleted relative to expectations given genomic GC content, with particularly strong depletion in mammals and many plant species but weaker depletion in monocots and some seedless plants (Fig. S6). Additionally, all four TpA-containing 3-mers are also consistently depleted, which is not true for 3-mers that contain ApT but not TpA dinucleotides (Fig. S6). These observations are consistent with prior reports of CpG and TpA dinucleotide depletion across distant eukaryotic and prokaryotic species (1, 2). In contrast, AAA/TTT were generally the most abundant 3-mers and enriched relative to expectations based on AT content of intergenic regions (Figs. S5 and S6); this enrichment is not driven by long A/T homopolymer tracts, which were removed as simple repeats.

Beyond these shared features, the trinucleotide composition varied substantially across species and showed clear taxonomic structure (Fig. 1A). Mammals had relatively similar profiles to one another and were distinct from plants, including lower relative abundance of CpG-containing 3-mers and CAA/TTG and higher abundance of CAG/CTG. Among plants, species within the same major clades tended to show more similar trinucleotide compositions (Fig. 1A; Fig. S5). Notably, members of the Poaceae family exhibited a distinctive profile characterized by relatively lower frequencies of AT-rich 3-mers and higher frequencies of GC-rich 3-mers, partially resembling the mammalian profiles. Unlike mammals, however, the Poaceae family showed substantially weaker depletion of CpG-containing 3-mers (Figs. S5 and S6).

Principal component analysis (PCA) revealed that most cross-species variation in intergenic trinucleotide composition was associated with two readily interpretable genomic features (Fig. 1B). Principal component 1 (PC1) explained 74.2% of the variance and was almost perfectly correlated with genomic GC content (Pearson’s r = −0.99; Fig. 1C), whereas PC2 explained an additional 11.4% variation and was strongly correlated with the CpG observed/expected (O/E) ratio (Pearson’s r = 0.93; Fig. 1D). Thus, GC content and CpG depletion correspond to the two dominant axes of variation in trinucleotide composition across the species examined. These observations motivated us to ask whether the observed trinucleotide compositions can be quantitatively explained as the equilibrium consequence of context-dependent mutation processes.

### A mutation spectrum-based Markov model predicts equilibrium trinucleotide composition

To predict the equilibrium trinucleotide composition associated with a given mutation spectrum, we developed an iterative discrete-time Markov-chain model (Fig. 2A; Methods). The model considers single-nucleotide substitutions whose rates depend on the substitution type and its immediate 5′ and 3′ flanking bases. For substitutions at the central position of a 3-mer, the transition rate is specified directly by the corresponding entry in the 96-channel mutation spectrum. For substitutions at either flanking position, the relevant mutation rate also depends on a neighboring base outside the focal 3-mer; we therefore calculate a composition-weighted average across the possible extended contexts. Because these weights depend on the current trinucleotide composition, the transition matrix is not fixed even under a given mutation spectrum. At each iteration, we construct the corresponding transition matrix, solve for its stationary trinucleotide distribution, and use that distribution to update the context weights for the next iteration until convergence (Fig. 2A; Methods).

**Figure 2.**
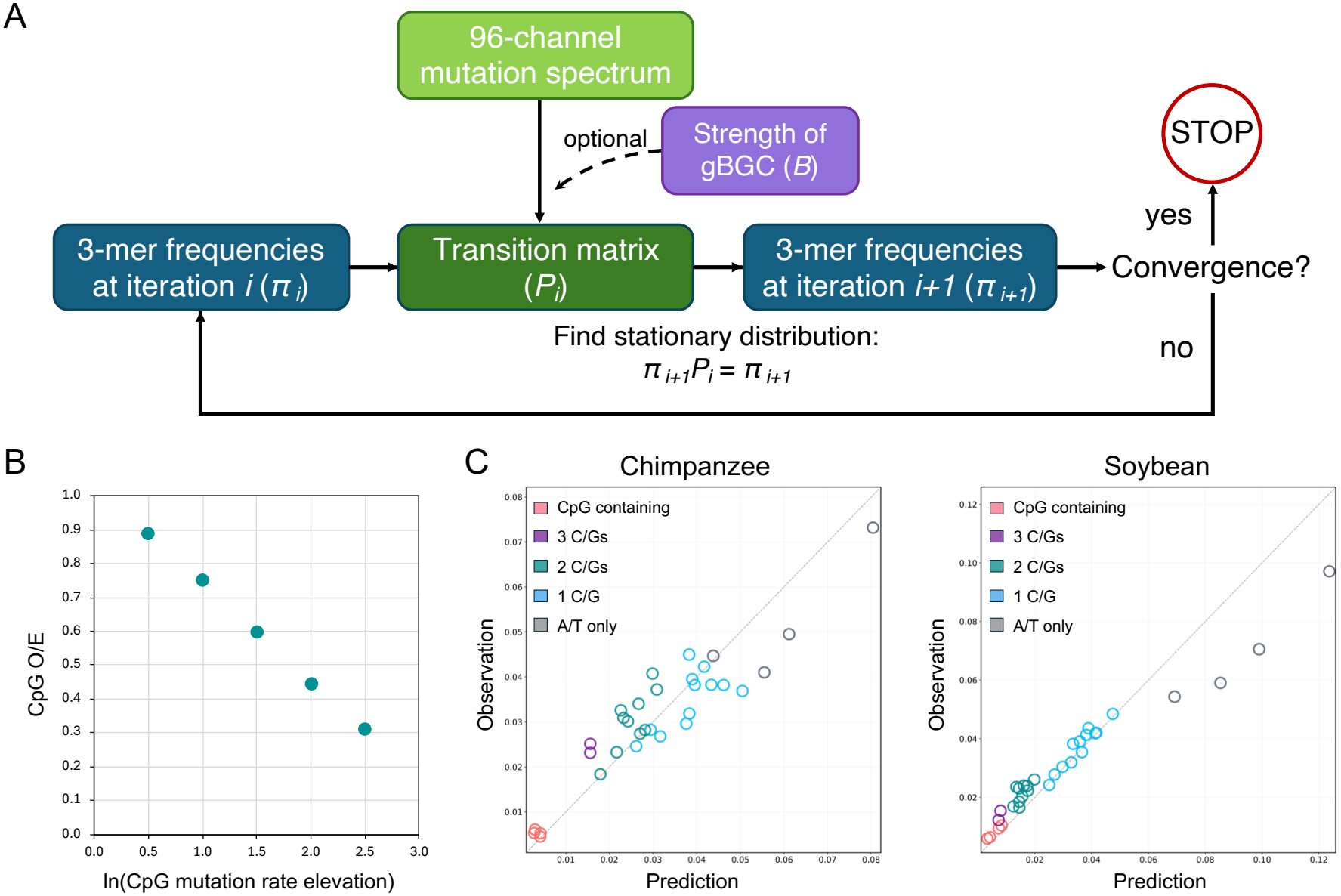
An iterative equilibrium model links a context-dependent mutation spectrum to trinucleotide genome composition. **A**. Schematic overview of the model and iteration workflow. At each iteration, the current 3-mer frequencies (*π*_i_) and a fixed 96-channel mutation spectrum are used to construct the transition matrix (Pi), which in turn defines the stationary 3-mer frequencies (*π*_i+1_) for the next iteration. This process is repeated until convergence. The model can optionally incorporate GC-biased gene conversion (gBGC) with strength (*B*). See Methods for more details. **B**. Application of the model to mock mutation spectra with progressively elevated CpG>TpG mutation rates recapitulates the expected progressive reduction in the CpG observed-to-expected (O/E) ratio at equilibrium. **C**. Comparisons between observed 3-mer frequencies in noncoding genomic regions and 3-mer proportions predicted by the iterative mutation-only model in chimpanzee and soybean. Each point represents one trinucleotide, with colors indicating GC content and CpG-containing status. The dashed line denotes the identity line.

As a positive control, we first examined whether the model could reproduce the expected compositional consequence of CpG hypermutability. We generated synthetic mutation spectra in which CpG transition rates were progressively increased while other mutation rates were held constant. Increasing CpG transition rates produced a monotonic decrease in the equilibrium CpG observed/expected ratio (Fig. 2B), closely mirroring the empirical finding based on polymorphism-derived mutation spectra (11).

Having established that the model captures this expected relationship between context-specific mutability and equilibrium composition, we next applied it to empirical mutation spectra and genomic compositions. Polymorphism-derived 96-channel mutation spectra were available for 25 plant and 10 mammalian species in our dataset (Fig. S7; Dataset S3)(11). Because these mutation spectra were inferred from noncoding variants, all subsequent comparisons between mutation spectra and genome composition were performed using matched noncoding sequences, including intergenic and genic noncoding regions (Dataset S4).

For each species, we used its polymorphism-derived mutation spectrum to predict the mutation-only equilibrium trinucleotide composition. The predicted and observed compositions showed substantial agreement in both mammals and plants, as illustrated by chimpanzee and soybean (Fig. 2C). Across all 35 species, the Jensen–Shannon divergence between predicted and observed 3-mer distributions was below 0.08 (Fig. S8; Table S1), indicating that contemporary context-dependent mutation spectra capture broad features of observed trinucleotide composition across diverse species. However, systematic deviations remained in most species (Fig. S8), pointing to contributions from fixation biases or other processes not captured by the mutation-only model.

### GC-biased gene conversion explains systematic deviations between mutation-only predictions and observed genomes

The residual deviations between predicted and observed compositions showed a consistent GC-directed pattern. Across most species, AT-rich 3-mers were overpredicted by the mutation-only model, whereas GC-rich 3-mers tended to be underpredicted (Fig. 2; Fig. S8). More generally, the equilibrium AT content predicted from mutation spectrum alone was high across all species (61–78%; Fig. 3A; Table S1), reflecting widespread mutation bias toward A/T (23). Yet the observed genomes were systematically more GC-rich than predicted from mutation alone (Fig. 3A), consistent with gBGC favoring the fixation of G/C alleles.

**Figure 3.**
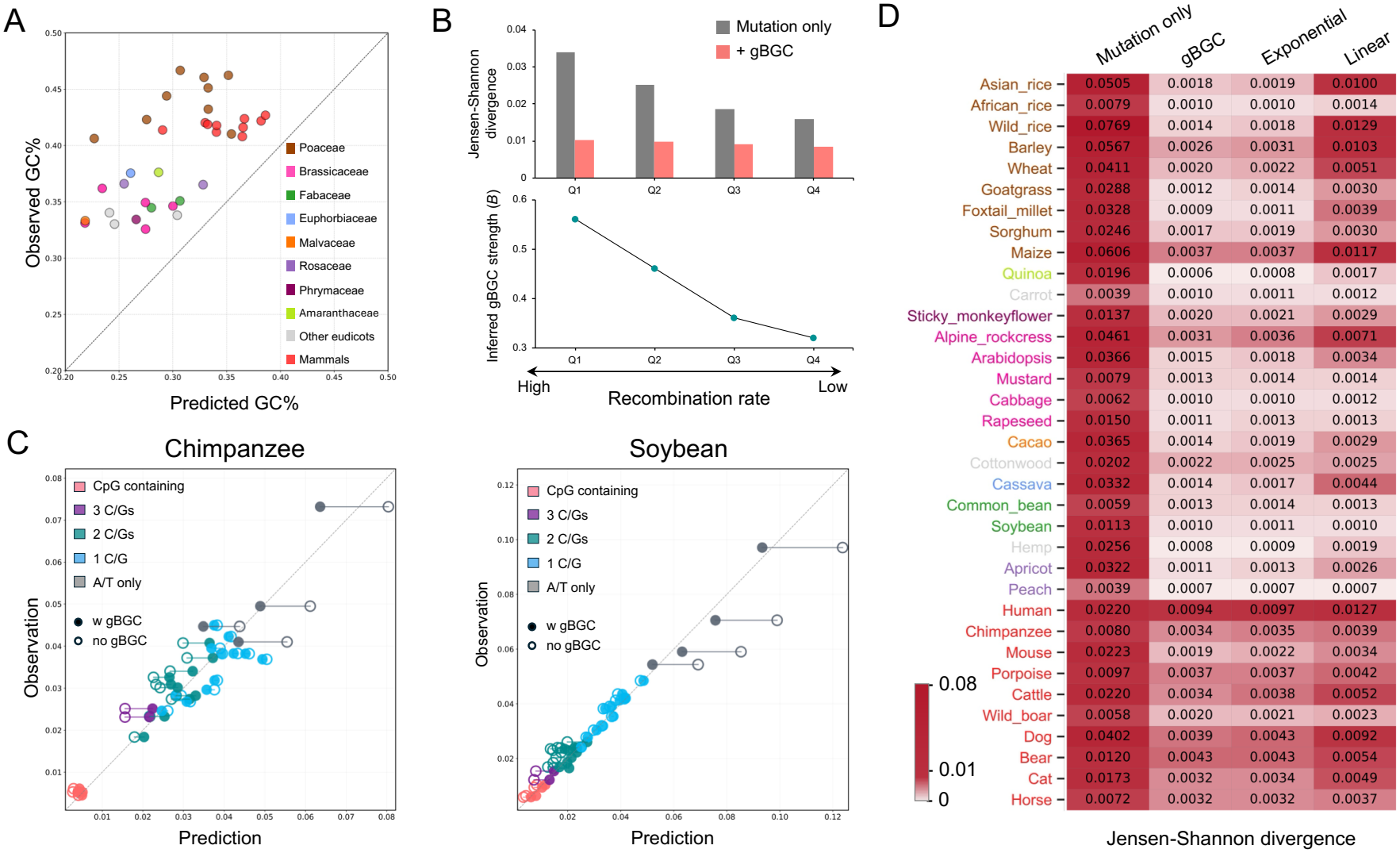
Incorporating the effects of gBGC greatly improves agreement between observed and predicted equilibrium trinucleotide genome composition. **A**. Comparison between observed GC% in noncoding regions and GC% predicted from the mutation spectrum across 25 plants and 10 mammals. **B**. Top, Jensen–Shannon divergence (JSD) between observed and predicted 3-mer frequencies under the mutation-only model across human genomic quartiles stratified by recombination rate, before and after incorporating gBGC. Bottom, inferred gBGC strength (*B*) required to match the predicted GC% to the observed GC% in each quartile. **C**. Comparisons between observed 3-mer frequencies and predictions with or without gBGC in chimpanzee and soybean. Each point represents one trinucleotide, with colors indicating GC content and CpG-containing status. Each horizontal solid line connects the predictions for the same trinucleotide under models with gBGC (filled) and without gBGC (hollow). The dashed line denotes the identity line. **D**. Jensen–Shannon divergence (JSD) between observed 3-mer frequencies and predictions under the mutation-only model, the gBGC model, an exponential GC-content correction, and a linear GC-content correction.

We next tested whether the magnitude of these deviations increased with recombination rate, as expected if gBGC contributed to the discrepancy. We divided the human autosomal genome into quartiles based on recombination rate and compared the observed composition of each quartile with the equilibrium predicted from the genome-wide mutation spectrum (Methods). The discrepancy between mutation-only predictions and observed composition increased with recombination rate (Fig. 3B), consistent with stronger gBGC in regions with higher recombination rates (19).

To determine whether a gBGC-like fixation bias could account for the remaining compositional differences, we extended the Markov model by modifying the relative fixation probabilities of AT- to-GC and GC-to-AT mutations according to the population-scaled strength of gBGC (*B*=4*N*_*e*_*b*) (Fig. 2A; Methods). Increasing B progressively increased the predicted frequencies of GC-rich 3-mers, including CpG-containing 3-mers, as well as overall equilibrium GC content (Fig. S9). For each species, we estimated an effective *B* by matching the GC content of the predicted equilibrium composition to that of the observed genome (Fig. S10). When the human genome quartiles were fitted separately, the best-fit B increased monotonically with recombination rate, and the residual deviations became more similar across quartiles (Fig. 3B), providing an independent check that the fitted parameter behaves as expected for gBGC.

Relative to the mutation-only model, incorporating gBGC reduced systematic deviation from the observation (Fig 3C; Fig. S8), reflected in substantially lower Jensen–Shannon divergence across all species, although the magnitude of improvement varied (Fig. 3D). For example, mouse and Asian wild rice showed substantially improved agreement after incorporating the fixation bias, whereas carrot, peach, wild boar, and horse showed only modest improvement. Surprisingly, human stood out as the species with the greatest residual divergence after incorporating gBGC, despite having by far the largest polymorphism dataset. One possible contributor is bias in the polymorphism-derived mutation spectrum used. Baymer does not explicitly account for recurrent mutation, which becomes increasingly relevant in very large samples (12, 30); additionally, singleton variants were excluded in the source study to reduce false positives (11). Both may distort the relationship between polymorphism probabilities and underlying mutation rates. Thus, residual discrepancies may reflect limitations in mutation-spectrum estimation or other unmodeled processes rather than failure of gBGC itself to contribute to genome composition.

Because the gBGC parameter *B* was fitted to match the observed genomic GC content, some improvement over the mutation-only model is expected simply from correcting the overall GC mismatch. We therefore asked whether explicitly modeling gBGC through mutation-specific fixation probabilities provides additional predictive value beyond a generic adjustment of GC content. We tested this by comparing the gBGC model with two simpler models that reweighted mutation-only predicted 3-mer frequencies according to their GC content, using either a linear or exponential adjustment, with all three models matching the observed genomic GC content (Methods). The gBGC model outperformed both the linear and exponential GC-content corrections in 33 of 35 species, with only marginally higher JSD than the best-fitting correction in porpoise and common bean. (Fig. 3D; Table S2). This result suggests that modeling gBGC through its effects on mutation-specific transition probabilities under a population genetic model captures additional structure in trinucleotide composition beyond a generic GC-content adjustment.

### Genome composition retains quantitative information about context-dependent mutation spectra

Given that mutation spectra predict broad features of genome composition, we examined how much sequence-context information is required for accurate prediction. We compared the full 96-channel trinucleotide mutation spectrum with simpler mutation models: the 6-channel model distinguishes only the six basic substitution types (C>A, C>G, C>T, T>A, T>C, and T>G), whereas the 7- and 9-channel models additionally treat CpG transitions and/or transversions as distinct mutation classes (Methods). For each model, we incorporated gBGC and fitted its strength to match the observed overall GC content.

Explicitly modeling CpG transitions improved predictions relative to the 6-channel model in all mammals and non-grass plant species, and adding separate CpG transversion classes produced further improvement in all mammals and 18 of 25 plant species (Fig. 4A; Table S3). With the exception of human, the full 96-channel model consistently produced the lowest Jensen– Shannon divergence across all species examined; in human, the 9-channel model performed slightly better, consistent with potential technical biases in the inferred mutation spectrum noted above. These results indicate that mutation rate heterogeneity across trinucleotide sequence contexts is important for accurately predicting trinucleotide composition.

**Figure 4.**
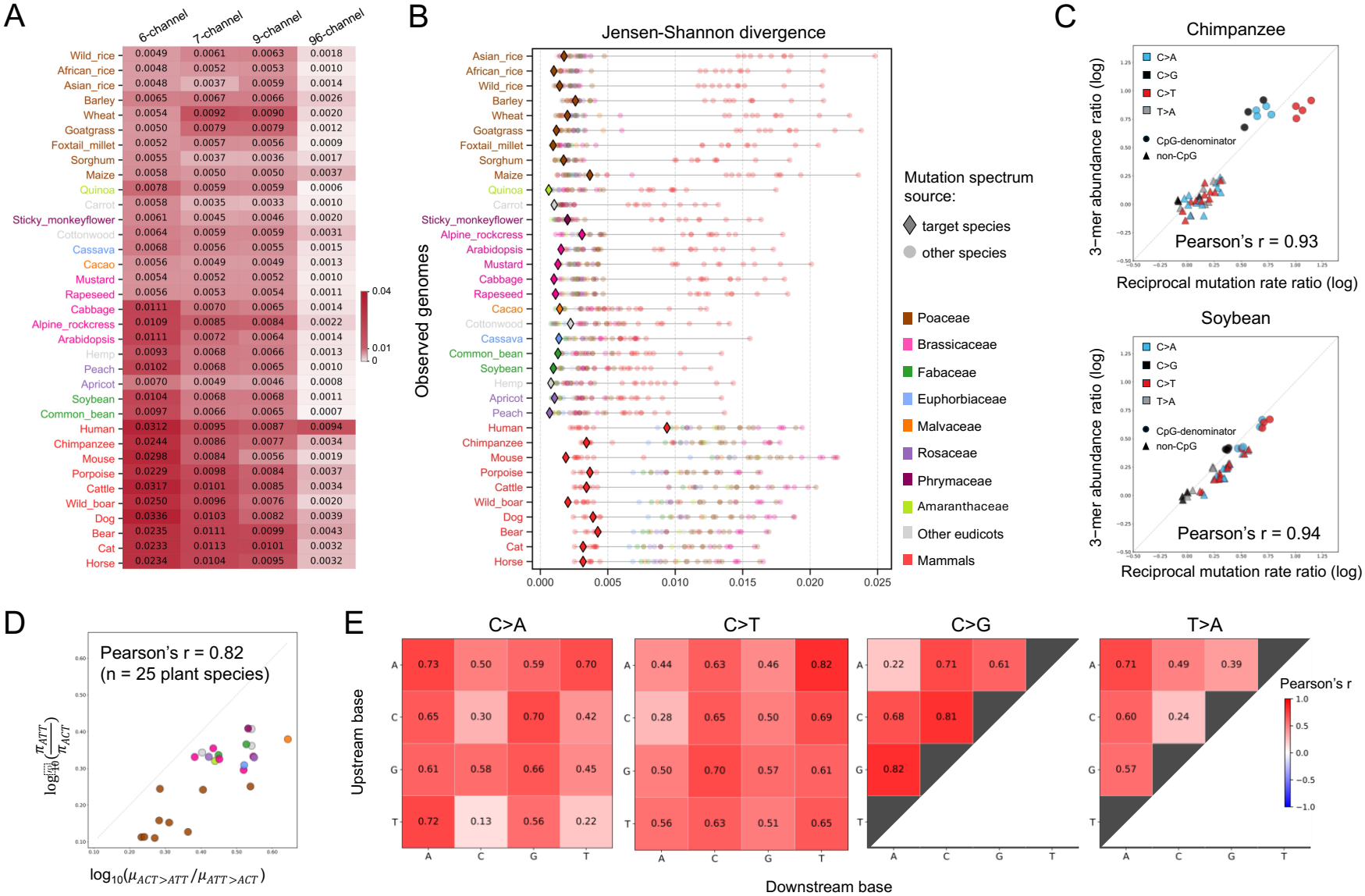
Genome composition reflects lineage-specific variation in context-dependent mutation spectra. **A**. Heatmap of Jensen–Shannon divergence (JSD) between observed 3-mer frequencies in noncoding regions and predictions using 6-channel, 7-channel, 9-channel, and 96-channel mutation spectra, with gBGC effects incorporated and strength adjusted to match the observed GC content. **B**. Comparison of JSD between observed and predicted 3-mer frequencies across species. Each row shows the JSD values between the observed genome composition of a given species and predictions using mutation spectra from each of the 35 species, with the gBGC strength independently adjusted for each genome–spectrum pair to match the observed GC content of the focal genome. Diamonds indicate predictions based on the focal species’ own mutation spectrum, whereas semi-transparent circles represent predictions based on mutation spectra from other species. Diamond and circle colors indicate the source species group of the mutation spectrum. **C**. Correlations between reciprocal mutation rate ratios and the corresponding 3-mer frequency ratios in chimpanzee and soybean. Both ratios were log10-transformed. All 44 possible pairs are shown. “CpG denominator” indicates pairs in which a CpG-containing 3-mer is used as the denominator in the 3-mer frequency ratio, for example, 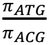 versus *μ*_*ACG > ATG*_*/μ*_*ATG*>*ACG*_. **D**. Correlation between the mutation rate ratio *μ*_*ACT*>*ATT*_*/μ*_*ATT*>*ACT*_ and the 3-mer frequency ratio 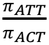 across 25 plant species. **E**. Heatmaps showing correlations between log10-transformed mutation rate ratios and 3-mer frequency ratios across 25 plant species. All 44 unique, non-reverse-complement trinucleotide pairs are shown, grouped by central-base mutation type, with upstream and downstream bases indicated on the y- and x-axes, respectively. Each cell shows the raw Pearson’s *r* value.

This observation raises the converse question of whether genome composition can distinguish among the context-dependent mutation spectra of different species. For each of the 35 species with a polymorphism-derived mutation spectrum available, we predicted its equilibrium trinucleotide composition using each of the 35 mutation spectra in turn and compared each prediction to the observed composition. We performed this analysis both under the mutation-only model and after incorporating gBGC, with the effective gBGC parameter *B* independently optimized for every genome–mutation spectrum combination to match the observed GC content. Thus, this analysis tests whether a species’ observed trinucleotide composition is preferentially predicted by its own mutation spectrum, and whether allowing variation in gBGC strength improves this correspondence.

Cross-species predictions showed strong taxonomic structure (Fig. 4B; Figs. S11 and S12). Plant genome compositions were generally predicted much better by plant mutation spectra than by mammalian spectra, and vice versa. Within plants, some finer-scale taxonomic structure was also apparent, most notably among grasses (Poaceae), whose mutation spectra tended to provide relatively good predictions for other grass species (Figs. S11 and S12). These patterns indicate that trinucleotide composition retains information about broad clade-specific differences in mutation spectra, even when gBGC strength is optimized separately for each comparison.

At the species level, however, the correspondence was considerably less specific. A species’ own mutation spectrum was often among the top predictors but was usually not uniquely the best-fitting spectrum (Fig. 4B; Fig. S12). Notably, allowing gBGC strength to vary generally improved the rank of a species’ own mutation spectrum relative to the corresponding mutation-only analysis (Fig. S12 C and D), indicating that accounting for variation in fixation bias strengthens the species-level correspondence between mutation spectra and genome composition. Nevertheless, the remaining limited resolution may partly reflect technical noise in mutation spectrum estimation and/or the long evolutionary timescale integrated by genome composition relative to contemporary polymorphism-derived mutation spectra. Thus, genome composition contains information about certain features of mutational processes but generally cannot identify a unique species-specific spectrum. Human was again a notable outlier: its own mutation spectrum predicted the observed human composition substantially more poorly than the other mammalian spectra and showed the poorest self-prediction among the species examined (Fig. 4B; Fig. S12C). This is again consistent with potential biases in the inferred human mutation spectrum. More generally, the occurrence of good fits among distantly related plant species (Fig. 4B; Fig. S12) indicates that the relationship between mutation spectrum and equilibrium trinucleotide composition is not one-to-one. This is also expected from their dimensionality: the 96-channel mutation spectrum contains substantially more independent dimensions than the 32-state trinucleotide composition, precluding unique recovery of the full mutation spectrum from trinucleotide frequencies alone. We thus turned to lower-dimensional summaries motivated by mutation equilibrium theory. In a two-state system, the equilibrium abundance ratio between two states is determined by the ratio of their reciprocal transition rates. Although trinucleotide equilibrium is more complex because each 3-mer is connected to multiple states, we hypothesized that the abundance ratios of reciprocal 3-mer pairs might nevertheless retain information about their relative mutation rates.

We first evaluated this relationship directly in the mutation-only Markov model. For 44 pairs of 3-mers differing by a single substitution at the central base, the equilibrium log-frequency ratio was strongly correlated with the corresponding log ratio of reciprocal mutation rates (Fig. S13; Pearson’s r ranging from 0.79 to 0.99; Table S4). Deviations from the identity line were expected because the equilibrium abundance of each 3-mer also depends on transitions involving its flanking positions and other connected states. Thus, despite the coupled nature of the trinucleotide system, reciprocal mutation-rate ratios capture a substantial component of the information underlying relative 3-mer abundance.

A similar relationship was detectable in observed genomes. Within individual species, log ratios of observed 3-mer frequencies were positively correlated with the corresponding reciprocal mutation-rate log ratios for most species, as illustrated by chimpanzee and soybean (Fig. 4C; Fig. S14; Pearson’s r ranging from 0.16 to 0.99 across all species; Table S4). These empirical correlations were generally weaker than those obtained from mutation-only equilibrium predictions, consistent with contributions from gBGC, error in mutation-spectrum estimation, and other evolutionary processes. Consistent with this interpretation, species inferred to experience stronger gBGC tended to show weaker correspondence between composition and reciprocal mutation rates (Fig. S15).

Finally, we examined whether the same relationship between 3-mer pair abundance and reciprocal mutation rates holds across species for individual reciprocal 3-mer pairs. For example, across the 25 plant species, the log ratio of ATT to ACT frequencies was strongly correlated with the log ratio of reciprocal mutation rates, log(*μ*_ACT_>ATT/*μ*_ATT_>ACT) (Pearson’s r=0.82; Fig. 4D). The observed abundance ratios were systematically shifted downward relative to the mutation rate ratios, in the direction expected under gBGC, illustrating how fixation bias can alter equilibrium abundances without completely eroding information about relative mutational input.

More broadly, all 44 reciprocal 3-mer pairs showed positive associations between cross-species variation in frequency ratios and mutation-rate ratios, although the strength of correlation varied among pairs (Fig. 4E; Figs. S16-S19; Pearson’s r ranging from 0.13 to 0.82; Table S5). Individual associations became weaker after accounting for phylogenetic relatedness, and none remained significant after multiple-testing correction (Methods). We therefore interpret these analyses not as evidence that individual mutation rate ratios can be inferred from genome composition, but as evidence that trinucleotide composition retains a detectable, lower-dimensional signature of lineage-specific mutation processes.

## Discussion

In the present study, we analyzed trinucleotide composition across 88 plant and mammalian genomes and found that most inter-species variation was associated with GC content and CpG depletion (Fig 1), two major features strongly shaped by mutation and fixation processes. To quantify these relationships, we developed a discrete-time Markov chain framework that predicts equilibrium trinucleotide composition from context-dependent mutation spectra and incorporates gBGC as a fixation bias. The mutation spectrum alone captured broad features of observed trinucleotide composition, and incorporating gBGC further reduced the systematic discrepancies between predicted and observed composition (Fig. 3D). The superior performance of the full 96-channel mutation spectrum over simplified mutation models further showed that fine-scale sequence-context dependence shapes the trinucleotide composition (Fig. 4A). We also demonstrated that genome composition retains partial information about lineage-specific mutation processes. Together, these findings provide a quantitative link between context-dependent mutation, fixation bias, and the long-term evolution of genome composition.

Interpreting this correspondence requires considering the different timescales represented by the mutation spectrum and genome composition. Genome composition integrates mutation and fixation processes over long evolutionary periods, whereas polymorphism-derived mutation spectra primarily reflect relatively recent processes. By connecting the two quantities, our analysis provides an indirect assessment of the stability of major features of mutation spectra and fixation biases: agreement is consistent with temporal persistence of these processes, whereas pronounced discrepancies may reflect bias or noise in the input data, model misspecification, or evolutionary non-equilibrium, including historical changes in mutation patterns or recombination landscapes.

Several sources of input-data uncertainty and model misspecification may limit concordance between model-based predictions and observed trinucleotide composition. Polymorphism-derived estimates of the mutation spectrum may be affected by sampling noise, recurrent mutation, and ancestral-state mis-polarization (31, 32). The genomic regions analyzed to minimize direct purifying selection may still contain unannotated functional sequences, and their definition depends on assembly and annotation quality (33, 34). The mutation model considers only single-nucleotide substitutions and their immediate sequence context, omitting extended-context effects and other types of mutational events, some of which may substantially alter composition. The framework also assumes spatially homogeneous mutation and fixation processes, despite evidence for regional heterogeneity in both (35). These limitations may contribute to residual discrepancies and currently limit our ability to distinguish evolutionary non-equilibrium from technical explanations.

Nevertheless, the close correspondence between predicted and observed trinucleotide composition across most species is consistent with broad temporal stability in some major features of mutation and fixation processes. In particular, the consistent excess of observed GC content over mutation-only predictions, together with the stronger discrepancy in high-recombination regions of the human genome, supports a widespread role for gBGC in opposing AT-biased mutational pressure (Fig. 3). Incorporating gBGC through a single parameter (*B*=4*N*_*e*_*b*) under a simplified population-genetic model improved predicted trinucleotide frequencies beyond generic GC-content corrections, outperforming the linear correction in all species and the exponential correction in most species (Fig. 3D). Notably, our inferred human value (B=0.42) closely matches previous estimates from allele frequency spectra (B=0.27–0.43) (31). This concordance between approaches using largely independent information suggests that genome composition can provide complementary information about effective genome-wide gBGC strength, although the inferred *B* may also absorb effects of unmodeled processes.

Perhaps most strikingly, genome composition also retains informative, although incomplete, summaries of lineage-specific mutation processes. Although its lower dimensionality precludes unique recovery of a full 96-channel mutation spectrum, trinucleotide composition still reflects broad taxonomic differences in mutation spectra and preserves quantitative information about relative mutation rates (Fig. 4B), as illustrated by the correspondence between reciprocal 3-mer abundance ratios and reciprocal mutation-rate ratios (Fig. 4E; Figs. 16-19). Notably, allowing gBGC strength to vary generally strengthened the correspondence between each species’ own mutation spectrum and its genome composition (Fig. S12), emphasizing the importance of accounting for fixation bias when interpreting these signals.

These properties suggest two complementary applications. In species lacking sufficient direct mutation data, genome composition may help generate hypotheses about mutational processes and identify unusual taxa for follow-up study. In species with richer data, it can provide a baseline for identifying discordance between contemporary mutational input and long-term genome composition. Extensions incorporating longer sequence context and regional heterogeneity could improve predictions and help distinguish model misspecification from genuine evolutionary non-equilibrium, enabling investigation of how mutation and fixation processes evolve across lineages and through time.

## Materials and Methods

### Data collection

We collected genome assemblies and corresponding gene annotations for a total of 88 species, including 75 plants and 13 mammals (Dataset S1). The plant dataset spans major lineages, including seedless plants, angiosperms, and gymnosperms, whereas the mammalian dataset includes representatives from Cetartiodactyla, Carnivora, Perissodactyla, Primates, Rodentia, and Proboscidea. For downstream analyses, chromosome-level sequences were retained where available, whereas all sequences of scaffold- or contig-level assemblies were used for species lacking chromosome-scale assemblies. For hemp, all scaffold sequences were included because a substantial proportion of the genome assembly remains unanchored. Sex chromosomes were excluded from all mammalian datasets. For the bear genome, because chromosome-level sequences are only available for the sex chromosomes, all remaining scaffold-level sequences were retained for downstream analyses. Sequence filtering and file reformatting were performed using SeqKit (v2.9.0)(36).

### Phylogenetic tree construction

To establish the evolutionary relationship for the plant dataset, a species tree was constructed using OrthoFinder (v3.0.1b1)(37) applied to protein sequences under default parameters. In addition to the 75 plant species shown in Fig. 1A, 13 additional species (Dataset S5) were included to improve the accuracy of phylogenetic inference. For most species with chromosome-level assemblies available, only genes located on chromosomes were included. For species with assemblies available only at the contig or scaffold level, all annotated genes were retained to maximize orthology recovery. For jointfir, pineapple, coconut, African rice, goatgrass, hemp, common hop, and melon, where chromosome-level assemblies are available but contained substantial numbers of unanchored genes, all genes were retained. For each gene, one representative transcript was retained: canonical transcripts were prioritized when designated in the species-specific GFF3 file; otherwise, the transcript with the longest coding sequence (CDS) was selected.

Phylogenetic inference was performed in two rounds using OrthoFinder. In the first round, protein sequences longer than 200 amino acids were used to identify orthogroups across all 88 plant species. Orthogroups were then filtered to retain those present in at least 90% of the species, containing no more than 15 gene copies per species, and with all member proteins at least 350 amino acids in length. For each retained orthogroup, only the longest protein sequence from each species was selected. These filtered protein sequences were then used as input for a second round of OrthoFinder analysis to infer the final plant species tree. The mammalian phylogeny was obtained from TimeTree (v5)(38) (accessed 05/2026). All phylogenetic trees were visualized and refined using iTOL (v6)(39).

### Simple repeat masking

Simple repeats and low-complexity regions were identified and masked using RepeatMasker (v4.1.7-p1)(40) with the following parameters: “-e rmblast -pa 32 -noint -gff”. All masked regions were subsequently excluded from downstream sequence extraction and composition analyses.

### Repetitive sequences masking strategy

For repetitive sequence masking, we compared four strategies in *Arabidopsis thaliana*, rice (*Oryza sativa*), and maize (*Zea mays*): (a) RepeatMasker using the Repbase repeat library (v31.05)(41); (b) RepeatMasker using the Repbase repeat library followed by additional low-complexity masking with DUSTMasker (v1.0.0)(42) using default parameters; (c) RepeatMasker using a *de novo* repeat library generated with RepeatModeler (v2.0.8)(43); and (d) species-specific published TE annotations or curated libraries. RepeatMasker was run with the same parameters for all analyses: -e rmblast -pa 6 -gff -q -no_is -norna -div 40 -cutoff 225. For maize, the B73v5 TE annotation was obtained from MaizeGDB (44). For Arabidopsis, transposable element annotations were obtained from the Araport11 annotation available through TAIR (https://www.arabidopsis.org/). For rice, the manually curated rice TE library (rice7.0.0.liban) was obtained from the riceTElib repository (45-47) and used with RepeatMasker to generate the corresponding masked genome.

### Definition of genomic compartments

For each genome with simple repeats masked, sequences were partitioned into three disjoint compartments: coding sequence (CDS), genic noncoding, and intergenic regions. Coding regions were defined based on CDS annotations in species-specific GFF3 files. Genic noncoding regions were defined as the union of introns and untranslated regions (UTRs). Intergenic regions were defined as all remaining genomic regions excluding both coding and genic noncoding sequences. Additionally, candidate regulatory regions were defined as the 1-kb regions upstream and downstream of each gene body (i.e., transcription start site to transcription end site).

For the human genome, conserved non-exonic regions were identified using the 92-mammal GERP constrained-element annotation from Ensembl release 116 (48). All annotated exonic regions from the NCBI RefSeq GRCh38.p14 annotation were excluded using BEDTools (49). The remaining regions represent non-exonic sequences under long-term evolutionary constraint, consistent with purifying selection.

For analysis of intergenic regions, we tested the effect of including and excluding candidate regulatory and conserved non-exonic regions in the human genome and found highly similar base compositions (Fig. S20). Therefore, we decided not to explicitly exclude potential regulatory non-coding sequences in base composition analysis across all species.

### Quantification of trinucleotide genome composition

To quantify trinucleotide (3-mer) frequencies across unmasked sequences of each genomic compartment, a sliding window of 3 bp with 1bp step size was applied. Windows containing ambiguous bases (e.g., ‘N’) or overlapping unmasked sequence boundaries were excluded. The observed proportion of each 3-mer was calculated as its count divided by the total number of observed 3-mers within the corresponding compartment. Counts for reverse-complementary 3-mer pairs (e.g., 5’-ATC-3’ and 5’-GAT-3’) were collapsed into 32 trinucleotide classes whose central base is a pyrimidine (C or T) to ensure strand-symmetrical composition profiling. This pipeline was implemented using custom bash/python scripts (see code availability below).

### Identification and length analysis of homopolymer tracts

A/T and C/G homopolymer tracts were identified on the forward strand using SeqKit (v2.9.0). The resulting intervals were converted to BED format, sorted, and merged using BEDTools (v2.30.0)(49), and the length of each merged tract was subsequently calculated.

### Calculation of observed/expected (O/E) ratios

The observed/expected (O/E) ratio for each 3-mer was defined as the ratio of its observed proportion to its expected proportion under a null model of independent nucleotide assortment, calculated based on compartment-specific base composition.

Single-nucleotide frequencies (P(*A*), P(*T*), P(*C*) and P(*G*)) were first estimated using the “stats” function in SeqKit (36). The expected frequency of a given 3-mer was then calculated as the product of the corresponding nucleotide frequencies. For example, for a trinucleotide *XYZ*, the expected frequency was calculated as:

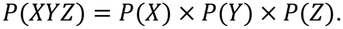

Similarly, the CpG O/E ratio for a genomic compartment was calculated as the observed proportion of CpG divided by the product of C and G proportions. The observed CpG frequency was determined using custom scripts similarly as quantification of 3-mer frequencies.

### Calibration of mutation spectra from polymorphism data

We downloaded the trinucleotide context-dependent relative mutation rates inferred by Ramos-Almodóvar et al., (2026) using Baymer (12), a Bayesian framework that estimates context– dependent polymorphism probabilities (*q*). Under low mutation rates and small sample sizes, the polymorphism probability at a given genomic site is approximately proportional to its per generation mutation rate (*μ*). However, this assumption breaks down as sample sizes increase and recurrent mutations become non-negligible, particularly in large human polymorphism datasets (50, 51).

To calibrate the per generation mutation rate (*μ*) from inferred polymorphism probability (*p*), we modeled their relationship using an exponential function:

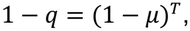

where *T* represents the total branch length of the coalescent tree connecting all sampled chromosomes, measured in generations. Solving for *μ* yields the transformation:

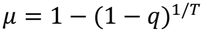

In this study, we set *T*=1,000,000 for all species. This value is arbitrary but has little effect on the inferred relative mutation rates for most species based on two rationales. First, when *q* is sufficiently small (*q*<0.05), the relative mutation rates (*μ*) across mutation types are approximately proportional to the polymorphism probabilities (*q*) across mutation types, largely independent of the exact value of *T*. Second, the predicted stationary composition depends primarily on relative transition probabilities rather than their absolute magnitudes (Fig. S21).

In addition, Almodóvar et al., (2026) excluded singleton variants (or variants only observed in one diploid individuals) to reduce false positives. This filtering can further distort the relationship between mutation rate and polymorphism probability, particularly in large samples where a substantial fraction of genuine recent mutations may be represented by singletons, and may therefore contribute to the poor correspondence between the human mutation spectrum and genome composition observed (Fig 3D; Fig. 4 A and B; Figs. S11 and S12). Because the effect of singleton exclusion depends on the absolute per-generation mutation rate, population demography, and sample size, we did not attempt to model this effect explicitly.

### Modeling stationary genome composition given a context-dependent mutation spectrum using a discrete-time Markov chain (DTMC)

Genome composition was modeled using a discrete-time Markov chain (DTMC). Because the mutation spectra used in our analysis were represented with reverse-complementary mutation types combined, we treated each trinucleotide and its reverse complement as a single equivalent state in the DTMC. The 64 possible trinucleotides were therefore reduced to 32 reverse-complement pairs. The stationary distribution vector *π* satisfies:

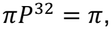

where *π* denotes the 1×32 vector of reverse-complement pair frequencies such that 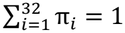, and *P*^32^ is the corresponding 32×32 transition probability.

We first constructed a 64×64 transition matrix *P*^64^ from the 96-channel mutation spectrum, based on the rules below. First, for transitions between states requiring two or more single-nucleotide substitutions (e.g., CCA→GAA), the transition probabilities were assumed to be zero. Second, for substitutions occurring at the central position of a trinucleotide (e.g., TTT → TAT), the transition probabilities were directly set to the corresponding relative mutation rates (*μ*_TTT → TAT_). For substitutions occurring at a side position of a trinucleotide, the transition probabilities were computed as context-weighted averages over all possible adjacent 3-mer configurations. For example, the transition probability from CCA to TCA (*μ*_CCA → TCA_) was defined as:

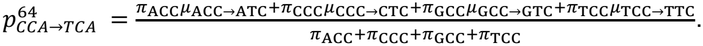

For each source state i, the self-transition probability on the diagonal was set to one minus the sum of all outgoing transition probabilities to other states in the same row:

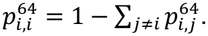

The resulting 64-state transition matrix *P*^64^ were then collapsed into a 32×32 transition matrix *P*^32^. For a source reverse-complement pair A = {a, RC} and a destination pair B = {b, RC}, the probability of transitioning from A to B was calculated as

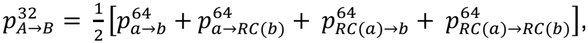

where RC represents the reverse-complement function. Note that two of the four terms above are necessarily zero, because the corresponding transitions involve two base substitutions.

Similarly, for each source reverse-complement state, the diagonal self-transition probability of *P*^32^ was calculated as one minus the sum of all outgoing transition probabilities to other reverse-complement states in the same row:

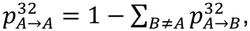

which is a faster way to calculate 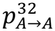 but nevertheless produces the same result by substituting B by A in the equation above.

The initial trinucleotide distribution was uniform across the 32 reverse-complement states, 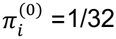. The stationary distribution *π* was computed using power iteration. At each iteration, *P*^(*k*)^ was re-calculated using the fixed mutation spectrum and *π*^(*k* − 1)^, the stationary distribution estimate from the previous iteration, until convergence was reached, defined as the sum of squared differences between successive estimates of *π* below 10^-8^:

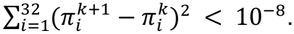

The converged stationary distribution was considered the predicted equilibrium trinucleotide composition.

### Simulation of synthetic mutation spectra to test CpG depletion

To evaluate the model’s ability to recapitulate CpG depletion in genomes with hypermutability at CpG sites (Fig. 2B), we generated a series of synthetic trinucleotide mutation spectra with pre-specified CpG hypermutability. In these experiments, the baseline mutation rates for all non-CpG 3-mer substitutions were fixed at 10^−8^. For CpG transitions (NCG>NTG), the mutation rates were scaled up by *e*^1/2^, *e, e*^3/2^, *e*^2^, and *e*^5/2^-fold, respectively, relative to the baseline rate, where *e* denotes Euler’s number. These synthetic mutation spectra were then passed to the DTMC model to predict the corresponding equilibrium trinucleotide compositions. CpG O/E ratios were subsequently calculated from the stationary compositions using the procedure described above.

### Comparing two trinucleotide genomic compositions using Jensen-Shannon divergence

The similarity between two trinucleotide distributions, either predicted versus observed or between two observed distributions, was quantified using the Jensen–Shannon divergence (JSD). Let *Y* and *Z* denote the two normalized trinucleotide frequency distributions, where fractions of all trinucleotides summed to 1. The Jensen–Shannon divergence (52) is defined as:

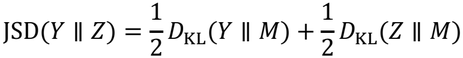

where 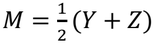 is the average distribution, and *D*_KL_ represents the Kullback–Leibler divergence (53) defined as:

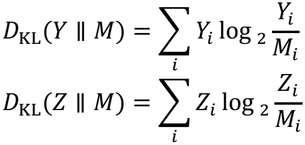

The logarithm was calculated using base 2, such that the resulting JSD values ranged from 0 to 1. A lower JSD value indicates greater similarity between the predicted and observed trinucleotide distributions, whereas higher values indicate greater divergence. Unlike standard Kullback-Leibler divergence, JSD is symmetric regarding *Y* and *Z*, always yields a finite value, and remains well-defined when specific trinucleotide frequencies are zero. These properties make JSD particularly suited for comparing discrete genomic compositions across species.

### Modeling GC-biased gene conversion (gBGC)

To model the effects of GC-biased gene conversion (gBGC) in our DTMC framework, we modified the transition matrix between 32 trinucleotide states by multiplying the mutation rate-based transition rates by the relative fixation probabilities due to gBGC, which impacts GC-altering mutation types (i.e., AT→GC or GC→AT mutations).

Specifically, gBGC was modeled as a fixation bias favoring AT→GC mutations over GC→AT mutations under a Wright–Fisher framework. The fixation probability of an AT→GC mutation was calculated as (20):

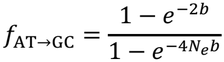

where *N*_*e*_ is the effective population size and *b* is the gBGC bias coefficient.

Similarly, the fixation probability of a GC→AT mutation was calculated as:

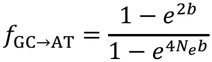

In contrast, the fixation probability for a GC-conservative (neutral) mutation is:

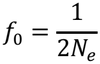

To model the relative fixation bias caused by gBGC, the above absolute fixation probabilities of different mutation types were normalized by *f*_0_:

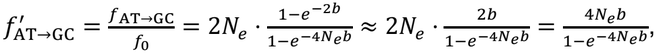

when *b* is small, as supported by current estimates in yeast and human (22, 54-57).

Similarly,

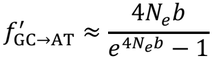

Redefining the population-scaled gBGC strength as *B* = 4*N*_*e*_*b*, the relative fixation probabilities for GC-altering mutation types can be expressed as:

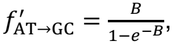

and

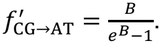

By assuming equal strength of gBGC bias on all GC-altering mutations, this model introduces only one additional parameter into our DTMC framework.

### gBGC strength estimation

We estimated *B*, the population-scaled gBGC strength, by minimizing the absolute difference in GC content between predicted and observed genome compositions. For each species, we first constructed a baseline transition matrix from the polymorphism-derived mutation spectrum. We then incorporated the directional bias of gBGC by multiplying the baseline transition rates by the relative fixation probabilities of GC-altering mutations, as defined above. The optimal *B* value was identified via a grid search across candidate values ranging from 0.00 to 1.00 with a step size of 0.01.

### Stratification of the human genome by local recombination rate

To evaluate the impact of local recombination rate and gBGC in explaining observed deviation between predicted and observed genomic composition, we performed analysis on regions of the human genome stratified by recombination rate. We first downloaded the HapMap GRCh38 genetic map available from the Beagle repository (https://bochet.gcc.biostat.washington.edu/beagle/genetic_maps/). The autosomal sequences (GRCh38) were initially partitioned into consecutive 10-kb intervals. Within each interval, boundaries were adjusted to align with the nearest flanking genetic markers from the HapMap GRCh38 genetic map. For each resulting interval, the local recombination rate was calculated as the ratio of genetic distance (*ΔR*) to physical distance (*ΔD*). Genomic intervals were then ranked by their local recombination rates (*ΔR*/*ΔD*) and stratified into quartiles, from Q1 (highest) to Q4 (lowest). Only autosomal sequences were included in this analysis.

### Exploring interactions between gBGC and different mutation types

For each mutation spectrum, we used the 32 reverse-complement trimer-pair proportions predicted by the DTMC model as the baseline distribution. To explore the effect of GC bias on this distribution, we compared two weighting strategies. For trimer pair i, with baseline proportion *p*_*i*_ and *k*_*i*_ G or C bases, the exponential strategy assigned 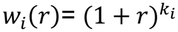 ,whereas the linear strategy assigned *w*_*i*_(r) = 1+*k*_*i*_*r*. Under either strategy, the adjusted proportion was *q*_*i*_(*r*) = *p*_*i*_*w*_*i*_(*r*)/∑_*j*_ *p*_*j*_*w*_*j*_(*r*), then the predicted GC% was calculated as ∑_*i*_ *q*_*i*_(*r*)*k*_*i*_*/*3. We evaluated r from 0 to 4 in increments of 0.01 and selected the value that minimized the absolute difference between predicted and observed GC%.

### Assessing context effects on mutation rates

To assess how sequence context affects mutation rates, we compared the original 96-channel mutation spectrum with three progressively simplified representations of each spectrum. First, we collapsed the 96 channels into six substitution classes by assigning each class its mean rate across all 16 trinucleotide contexts. Second, we created a 7-channel representation by separating C>T mutations at CpG sites from C>T mutations in other contexts. Third, we created a 9-channel representation by separating CpG and non-CpG mutations for each of C>A, C>G, and C>T. We compared results from the 96-, 6-, 7-, and 9-channel representations to evaluate the contribution of overall trinucleotide context and, more specifically, CpG context. Averaging was performed separately for each input spectrum.

### PCA for genomic trinucleotide composition

Principal component analysis (PCA) was performed on the 3-mer genomic composition profiles of intergenic regions of 82 species (Fig. 1B) using the “sklearn.decomposition.PCA” function implemented in scikit-learn (v1.6.1)(58), with “n_components=None” to retain all principal components.

### Phylogenetic regression

The phylogenetic regression analyses were performed to examine the relationship between 3-mer pairs abundance and reciprocal mutation rates ratios across 25 plant species using the “phylolm” R package (59), with the species tree inferred by OrthoFinder (37). A Brownian motion (BM) model of trait evolution was applied to account for phylogenetic relationships among species.

## Supporting information

Supplementary figures, legends for supplementary datasets

## Code availability

The code used for the present study is available at GitHub: https://github.com/yijie-wang11/TrimerCompositionAndMutationSpectrum.

## Author Contributions

Y.W. and Z.G. designed research; Y.W. performed research and analyzed data; Y.W. and Z.G. interpreted results; Y.W. wrote the original draft; Z.G. supervised the research and acquired funding; and Y.W. and Z.G. reviewed and edited the manuscript.

## Acknowledgments and funding sources

We thank Iain Mathieson, Ben Voight, Fabian Ramos-Almodóvar, and other members of the Gao and Mathieson laboratories for helpful discussions; and Fabian Ramos-Almodóvar for providing access to trinucleotide mutation spectra of 35 species prior to publication.

This work is supported by a Research Fellowship (FG-2021-15702) from the Alfred P. Sloan Foundation (https://sloan.org/) and a grant (R35GM146810) from the National Institute of General Medical Sciences to ZG.

