## Supplementary figures, legends for supplementary datasets for "Genomic trinucleotide composition reflects context-dependent mutation spectra in plants and mammals"

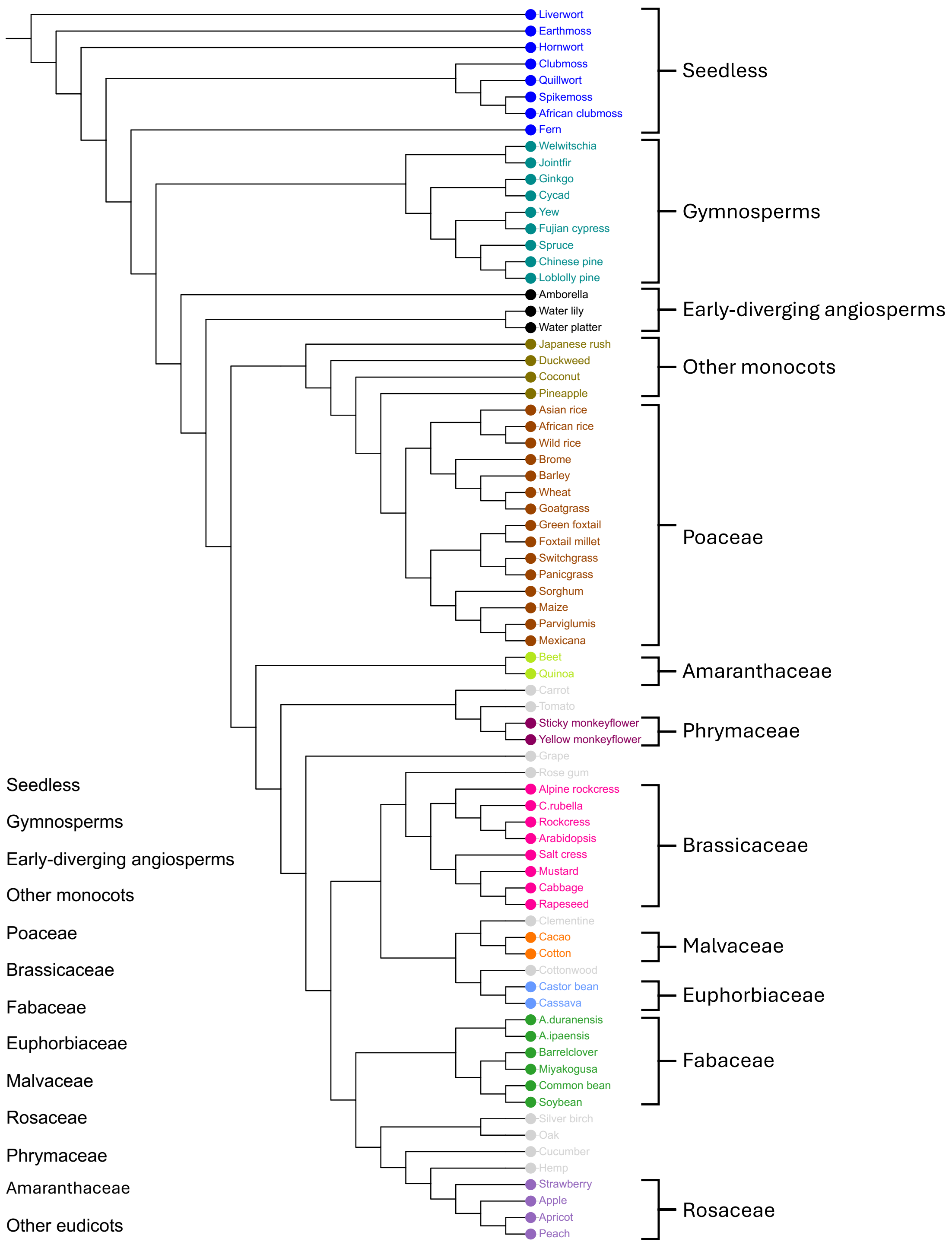


**Fig. S1 | Phylogenetic topology for 75 plant species.**
Only the topology of species relationships is shown; branch lengths do not represent genetic distance or divergence time. Leaf colors indicate species groups, as shown in the legend at the lower left.

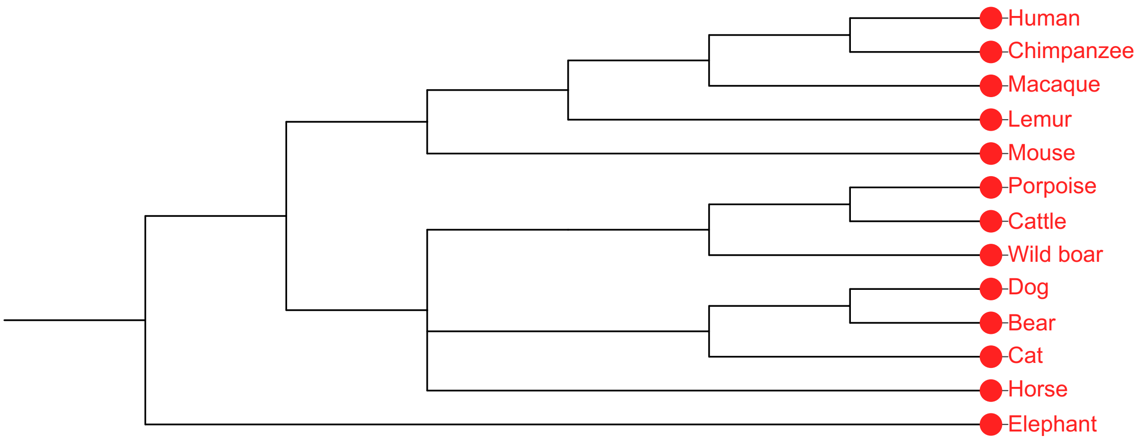


**Fig. S2 | Phylogenetic topology for 13 mammal species.**
Only the topology of species relationships is shown; branch lengths do not represent genetic distance or divergence time.

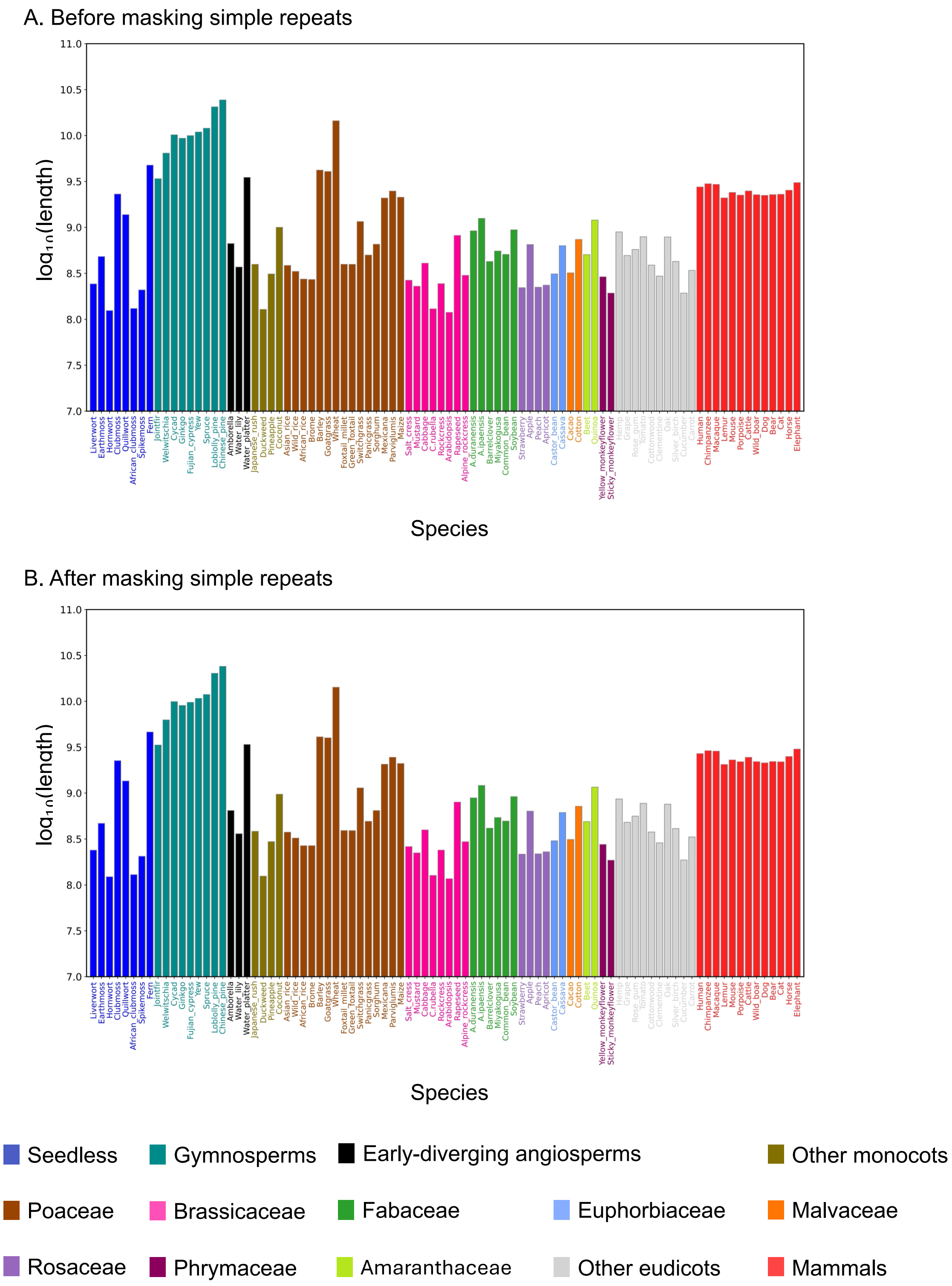


**Fig. S3 | Genome base counts across 88 species.**
Each bar represents the total number of genomic bases (A, T, C, and G) for each species before (**A**) and after (**B**) masking simple repeats with RepeatMasker. Base counts were $\log_{10}$-transformed

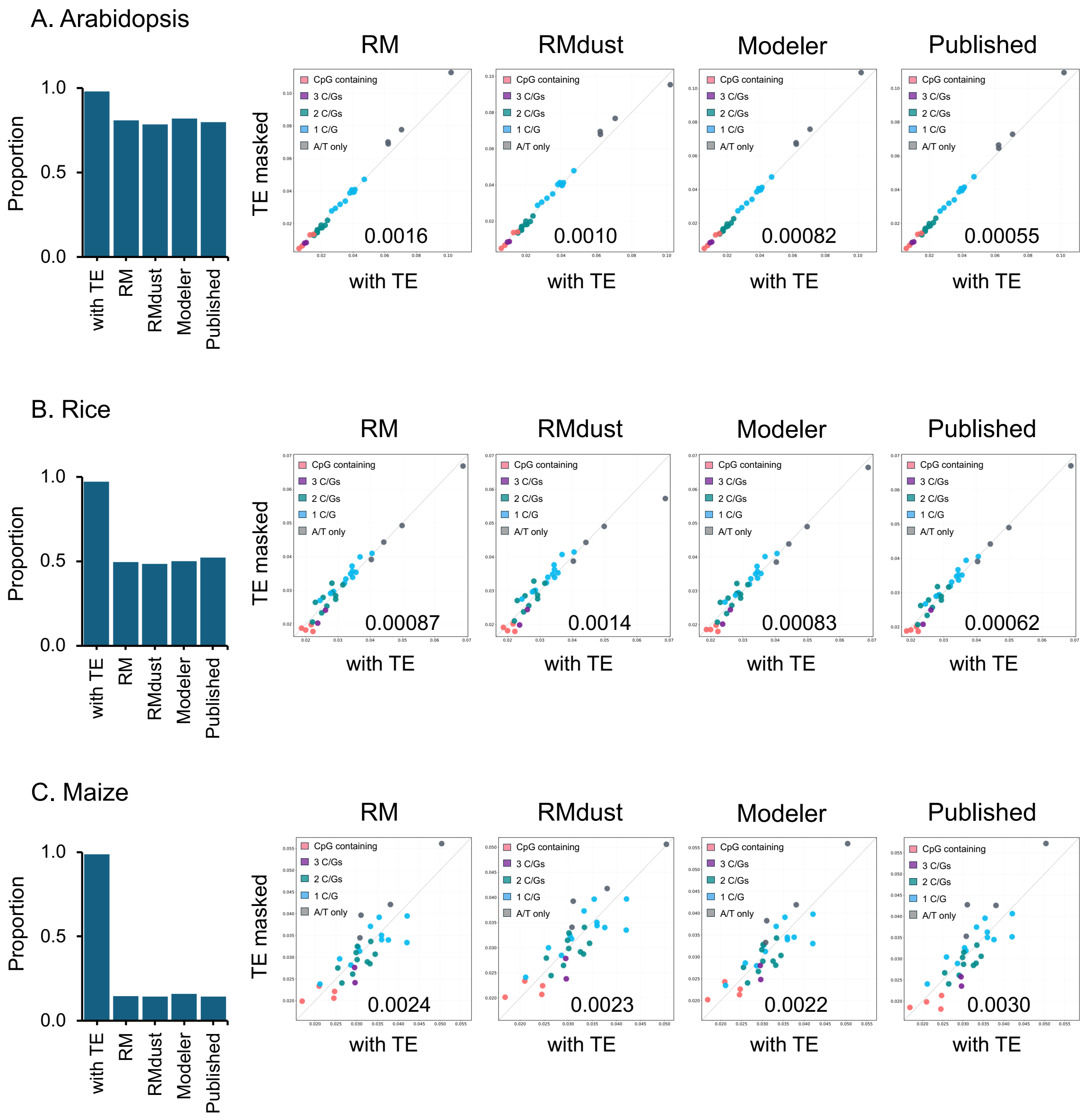


**Fig. S4 | Comparison of 3-mer frequencies across different repeat-masking strategies.** Panels show comparisons of 3-mer frequencies before and after TE masking in *Arabidopsis* (**A**), rice (**B**), and maize (**C**). For each species, the bar plot shows the proportion of the genome remaining unmasked under each strategy. "With TE" denotes only masking only simple repeats and low-complexity regions with RepeatMasker; "RM" denotes RepeatMasker using the Repbase repeat library; "RMdust" denotes RepeatMasker using the Repbase library followed by additional low-complexity masking with DUSTMasker; "Modeler" denotes RepeatMasker using a *de novo* repeat library generated with RepeatModeler; and "Published" denotes masking based on species-specific published TE annotations or curated repeat libraries. The scatter plots show 3-mer frequencies before TE masking ("With TE," x-axis) and after TE masking (y-axis) under each strategy. Point colors indicate 3-mer classes based on CpG presence and GC content. The Jensen–Shannon divergence for each comparison is indicated in the lower-right corner of the corresponding scatter plot.

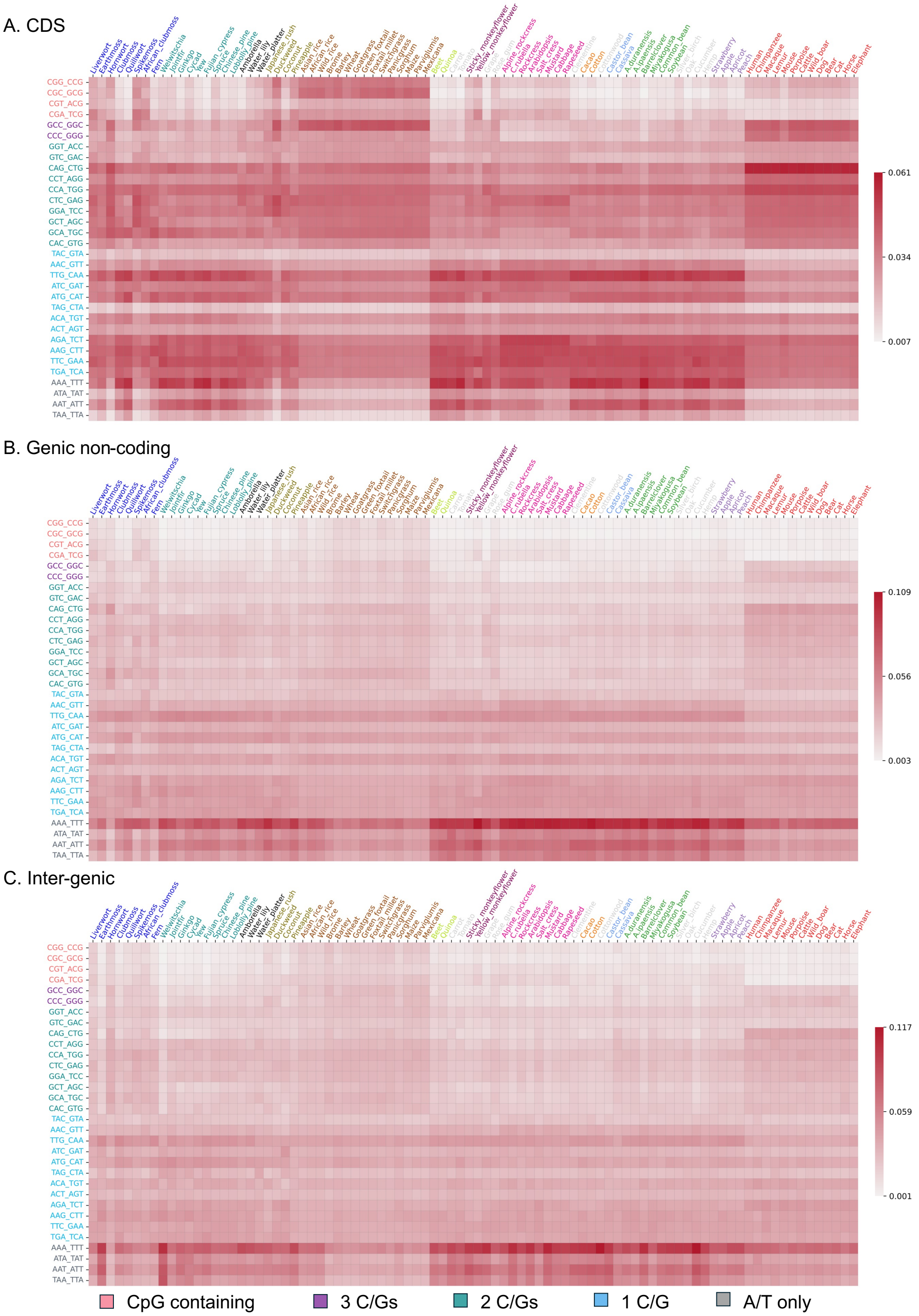


**Fig. S5 | Observed 3-mer frequencies across 88 species.**
Heatmaps show observed 3-mer frequencies after collapsing reverse-complement pairs in coding sequences (CDS) (**A**), genic noncoding regions (**B**), and intergenic regions (**C**). Species label colors indicate species groups, as in Figs. S1 and S2. 3-mer label colors indicate classes based on CpG presence and GC content (legend below panel C).

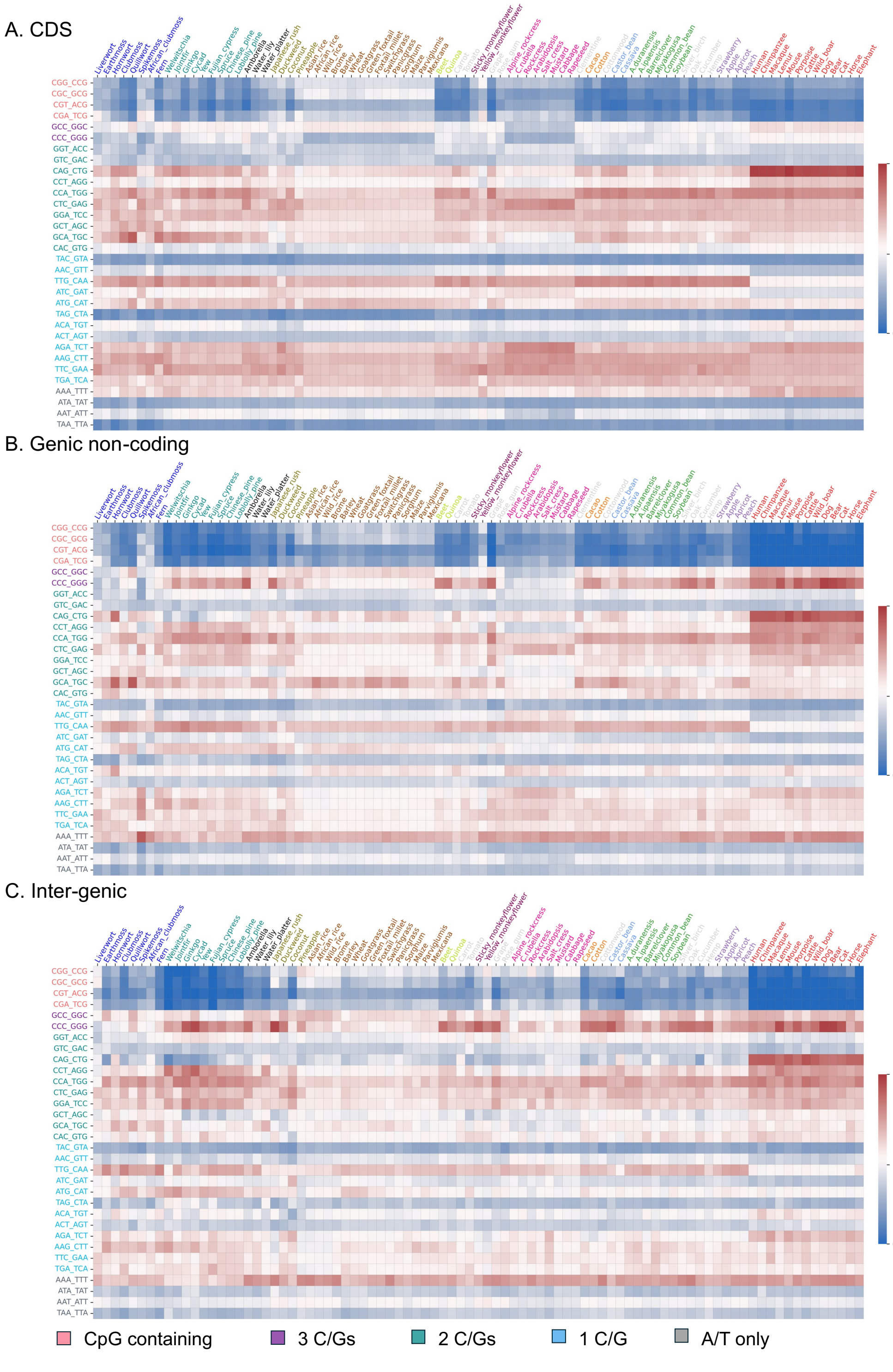


**Fig. S6 | 3-mer observed-to-expected ratios across 88 species.**
Heatmaps show observed/expected (O/E) ratios of all 3-mers in coding sequences (CDS) (**A**), genic noncoding regions (**B**), and intergenic regions (**C**). Expected proportions are based on compartment-specific GC content.

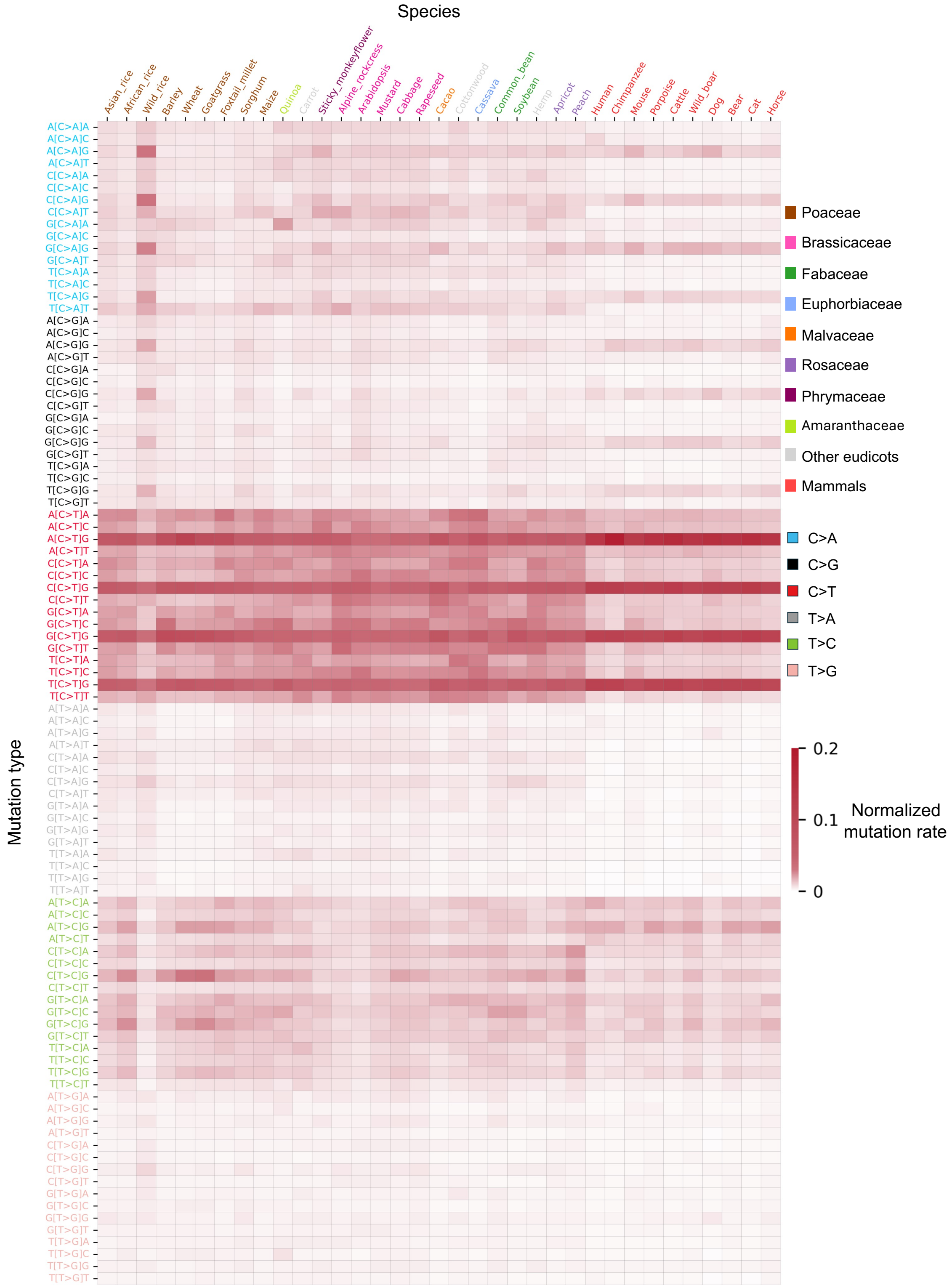


**Fig. S7 | Relative mutation rates across 35 species.**
Heatmaps show 96-channel mutation rates normalized within each species to sum to unity.

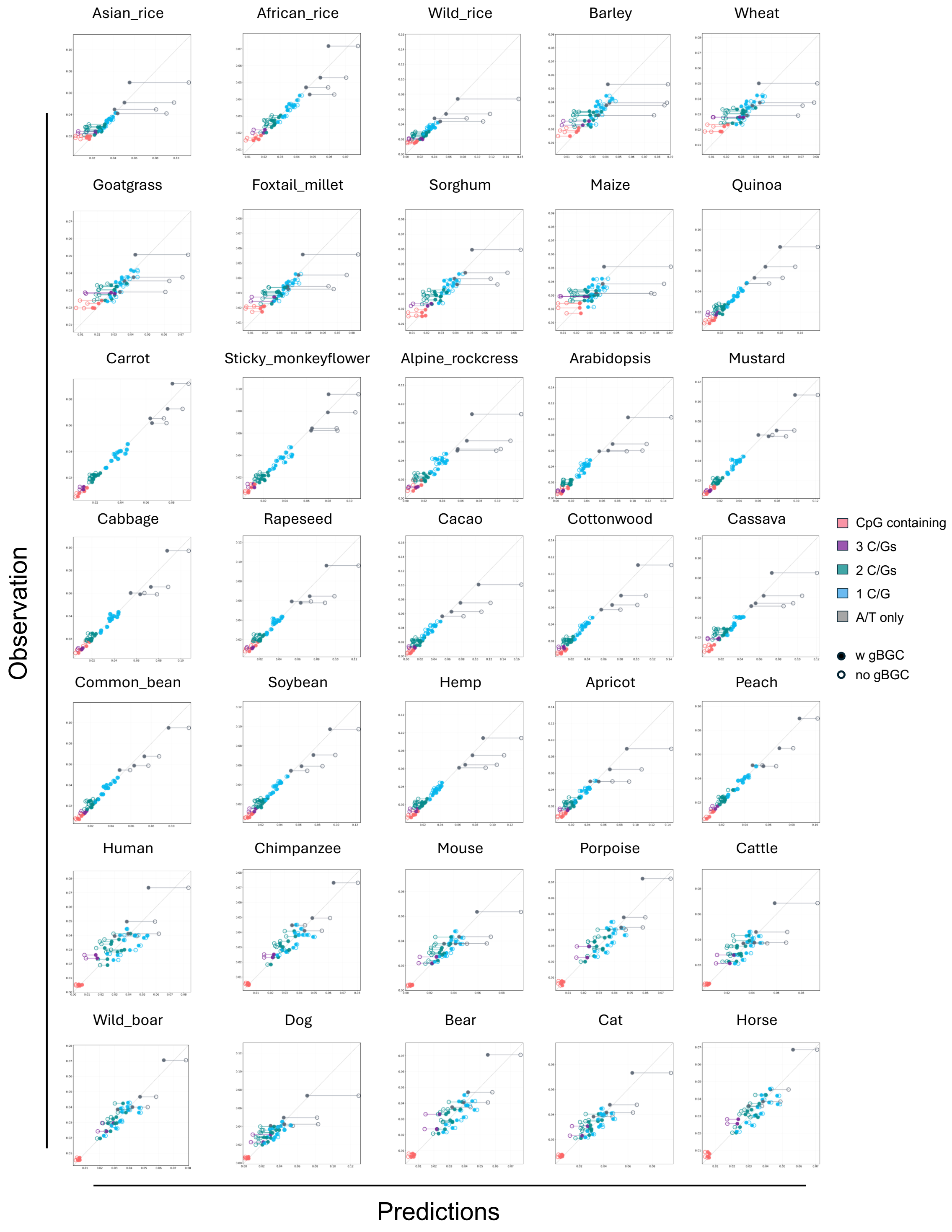


**Fig. S8 | Comparison between observed and predicted 3-mer frequencies in 35 species.** Scatter plots compare observed 3-mer proportions with proportions predicted with or without gBGC. The x-axis shows observed proportions, and the y-axis shows predicted proportions. For each 3-mer, a solid horizontal line connects the predictions with gBGC (filled point) and without gBGC (hollow point).

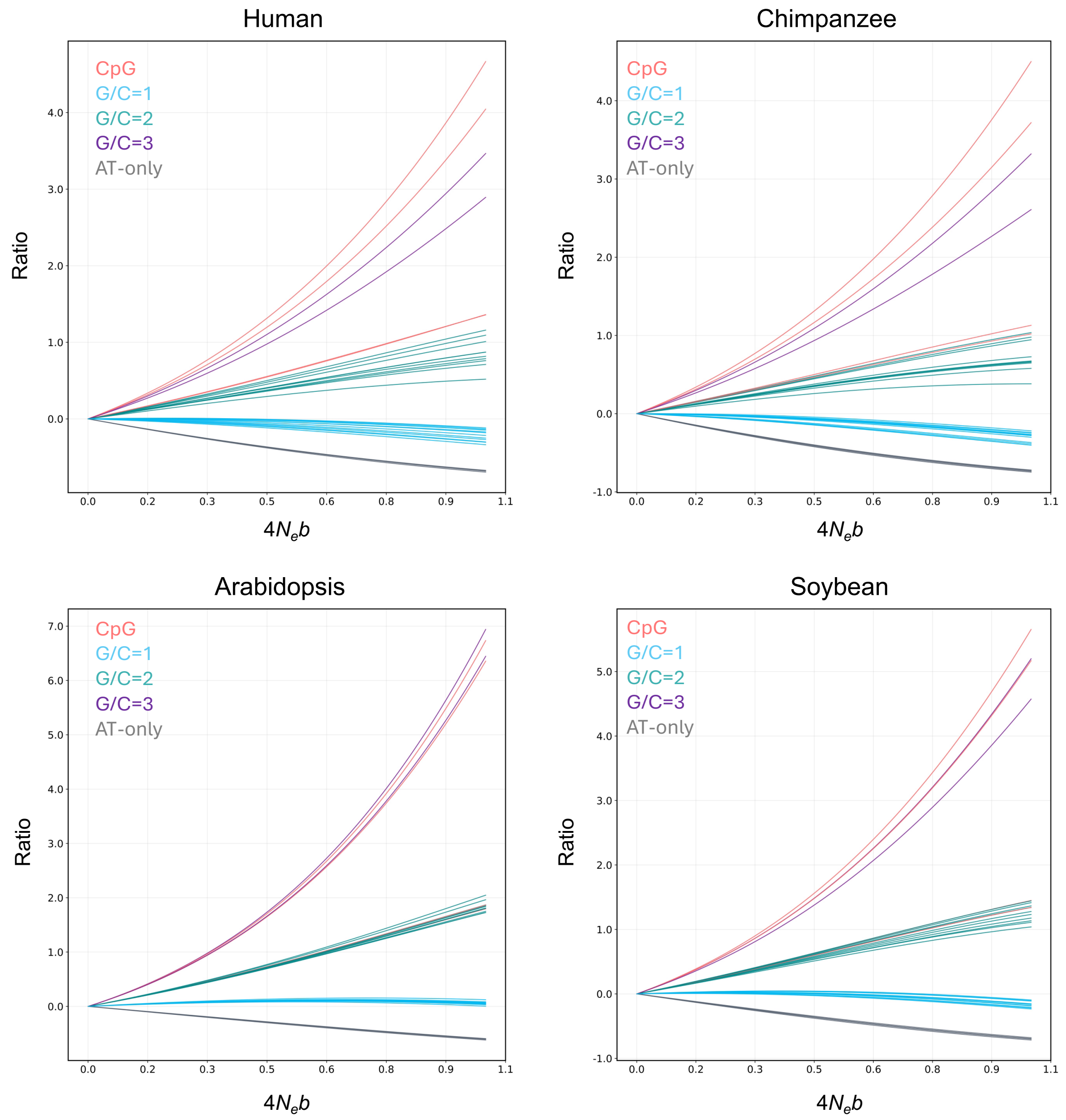


**Fig. S9 | Changes in equilibrium 3-mer frequencies under varying gBGC strengths (4*N*eb).**
Each curve represents one 3-mer after collapsing reverse-complement pairs. The x-axis shows gBGC strength ($4N_eb$) ranging from 0 to 1, measured as, ranging from 0 to 1. The y-axis shows the relative change in 3-mer frequency from its value at $4N_eb$ = 0, calculated as $(\pi_{4Neb} - \pi_0)/\pi_0$.

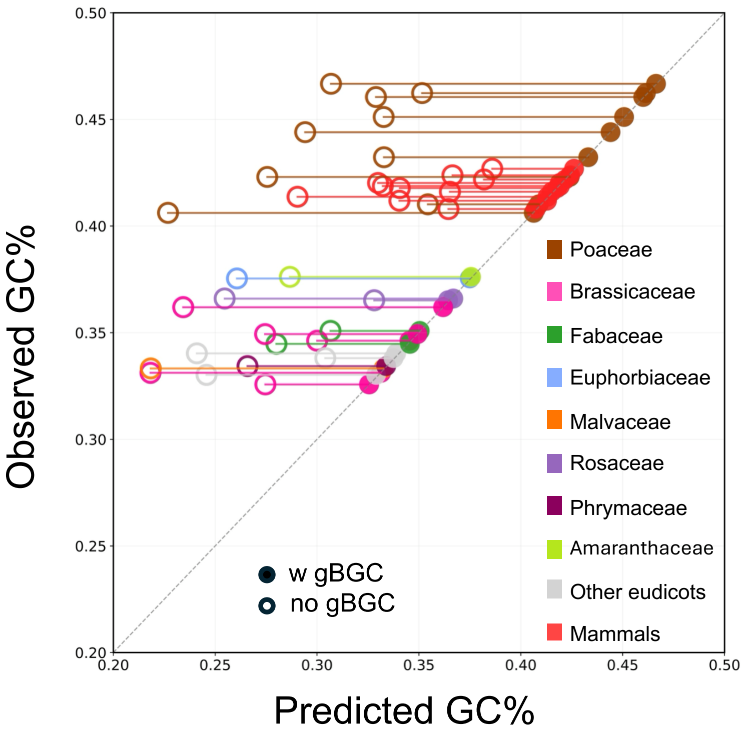


**Fig. S10 | Comparison between observed and predicted GC content with and without gBGC.** The scatter plot shows predicted GC content (x-axis) and observed GC content (y-axis) across 35 species. Predictions use each species' polymorphism-derived mutation spectrum. Solid lines connect paired predictions with gBGC (filled point) and without gBGC (hollow point). gBGC strength was adjusted to match each species' observed GC content.

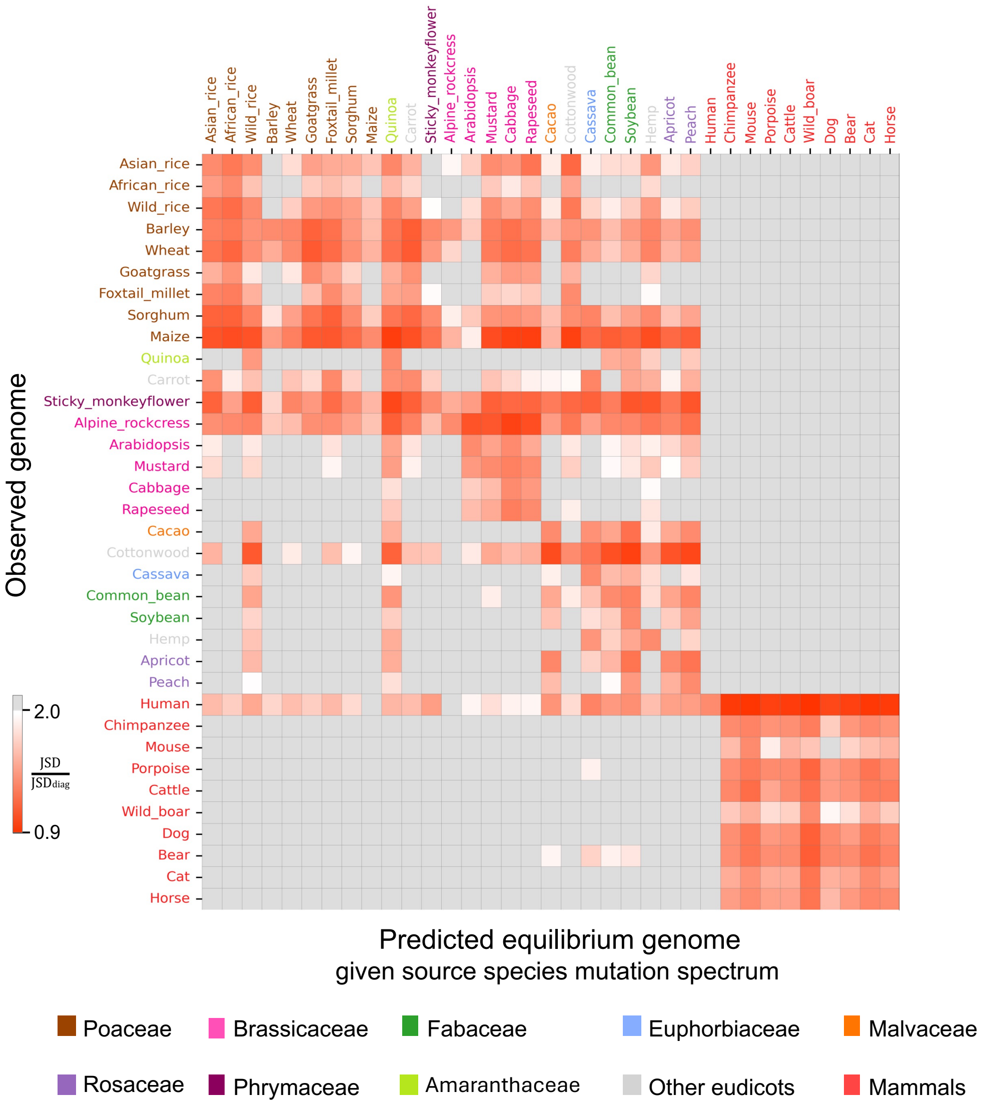


**Fig. S11 | Comparison of observed and predicted 3-mer composition across 35 species.**

Each row represents the observed 3-mer composition of a species, and each column indicates the species supplying the mutation spectrum. For each row–column pair, gBGC strength was adjusted to match the observed GC content of the row species. Each cell shows the Jensen–Shannon divergence (JSD) between the predicted equilibrium and observed compositions, normalized by the JSD of the same-species comparison in that row (the diagonal). Gray indicates normalized JSD values greater than 2.

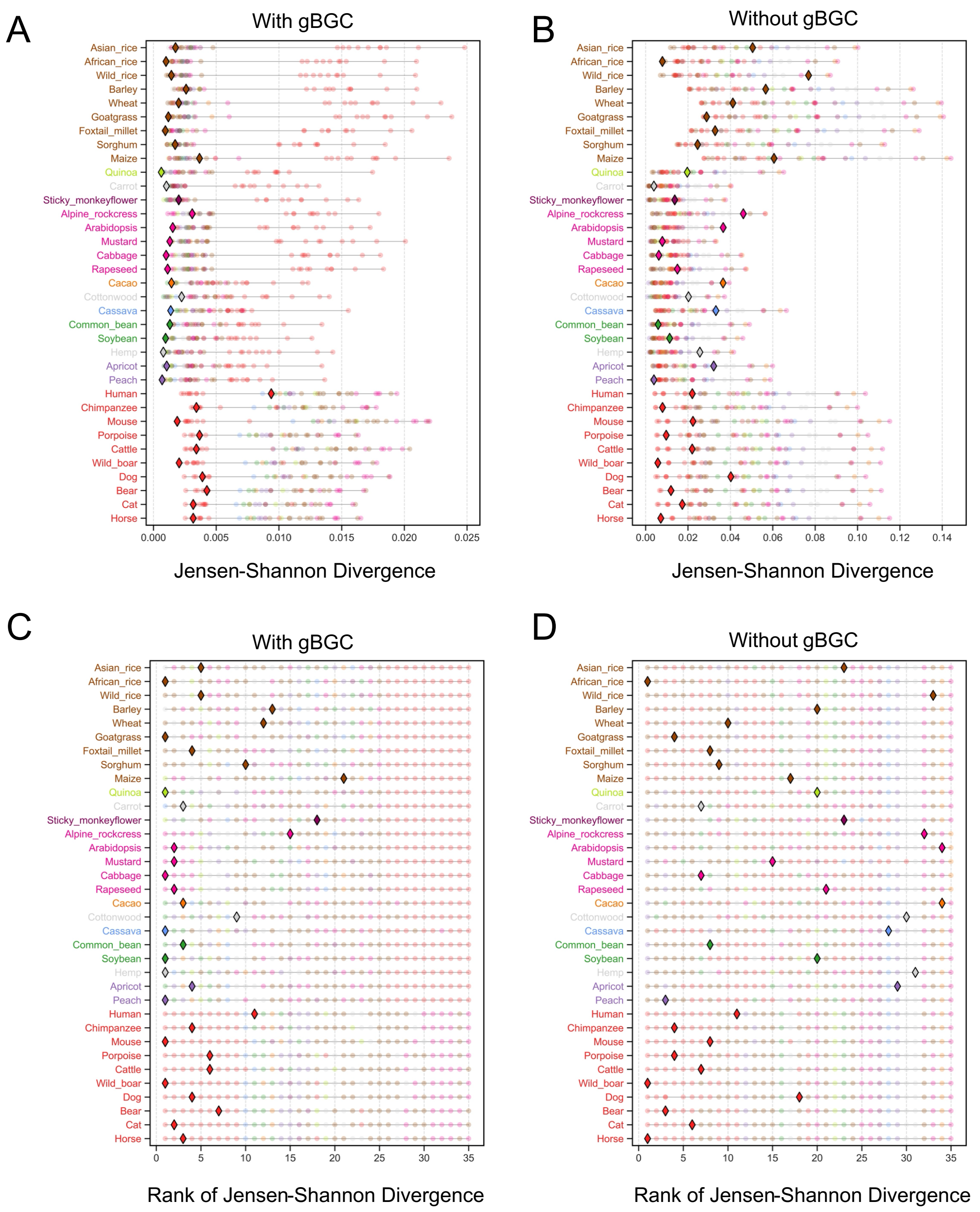


**Fig. S12 | Comparison of JSD between observed and predicted 3-mer compositions across species.**

**A–B.** JSD values between observed and predicted 3-mer proportions across species, with (**A**) and without (**B**) gBGC. Each row compares a focal species' observed composition with predictions based on each of the 35 species' mutation spectra. In (**A**), gBGC strength was adjusted separately for each comparison to match the focal species' observed GC content.

**C–D.** Within-row ranks of the JSD values in (**A**) and (**B**), respectively, with rank 1 indicating the lowest JSD.

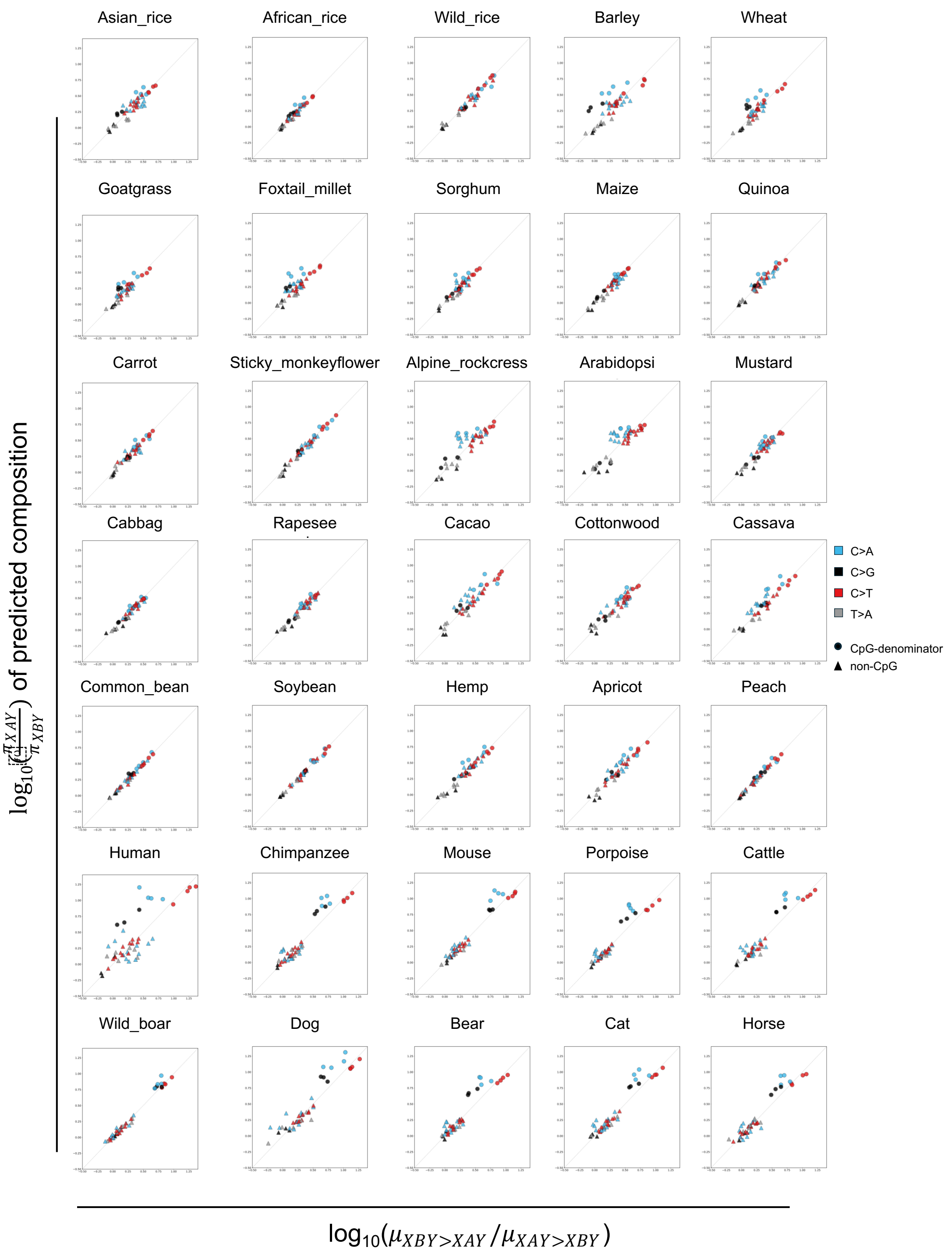


**Fig. S13 | Comparison of mutation rate ratios and predicted 3-mer proportion ratios across 35 species.**

For each species, the x-axis shows the ratio of reciprocal mutation rates between a pair of 3-mers that differ only at the central base, and the y-axis shows the corresponding ratio of equilibrium frequencies predicted without gBGC. Both ratios were $\log_{10}$-transformed.

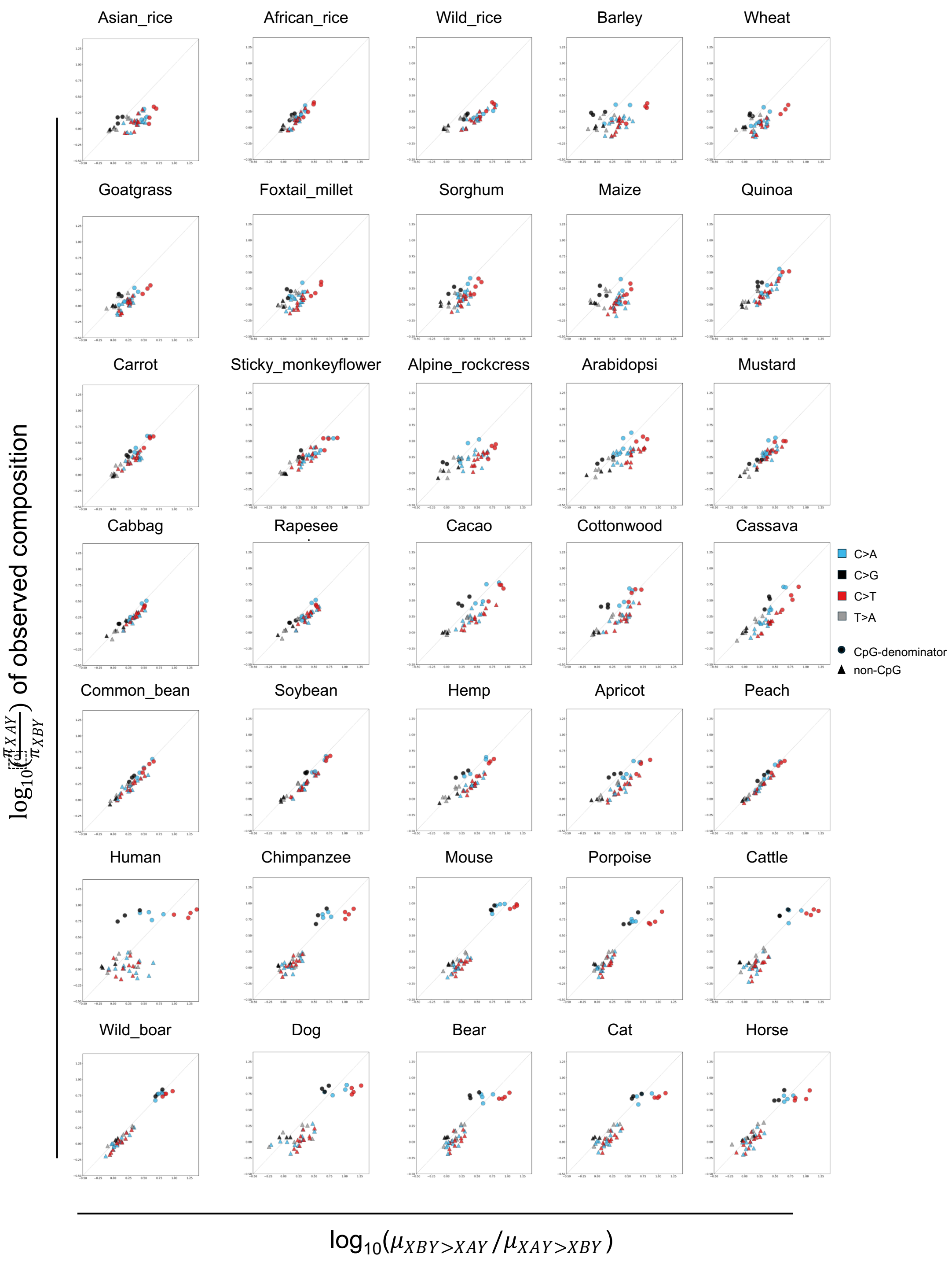


**Fig. S14 | Comparison of mutation rate ratios and observed 3-mer proportion ratios across 35 species.**
Same as Fig. S13, except that y-axis shows ratio of observed 3-mer frequencies.

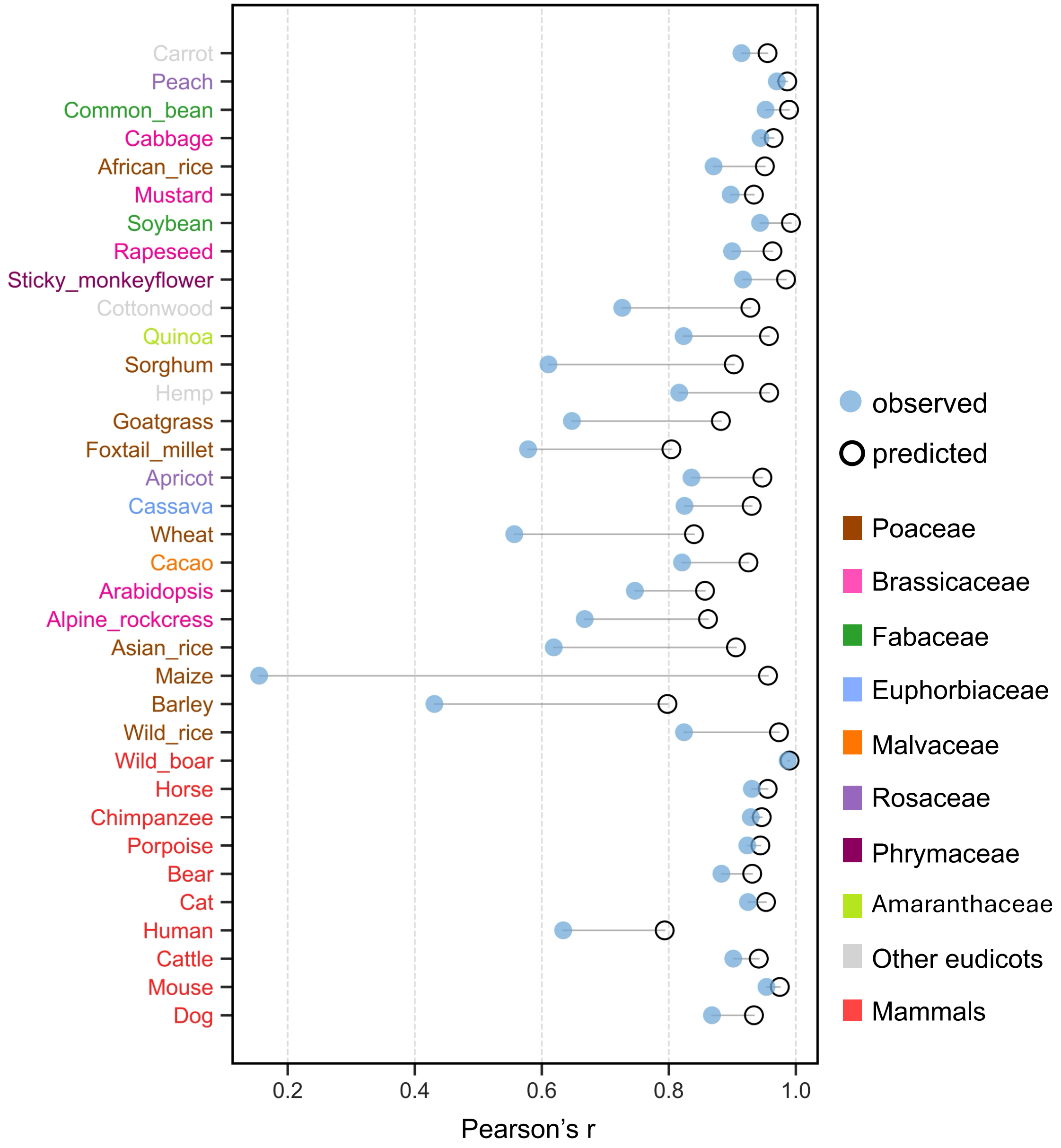


**Fig. S15 | Pearson's r between 3-mer proportion ratios and reciprocal mutation rate ratios across 35 species.**

Filled and hollow circles show correlations using observed 3-mer composition and those predicted with and without gBGC, respectively. Species are ordered by increasing inferred gBGC strength (top to bottom), separately within plants and mammals. Pearson's correlations were calculated on $\log_{10}$-transformed ratios.

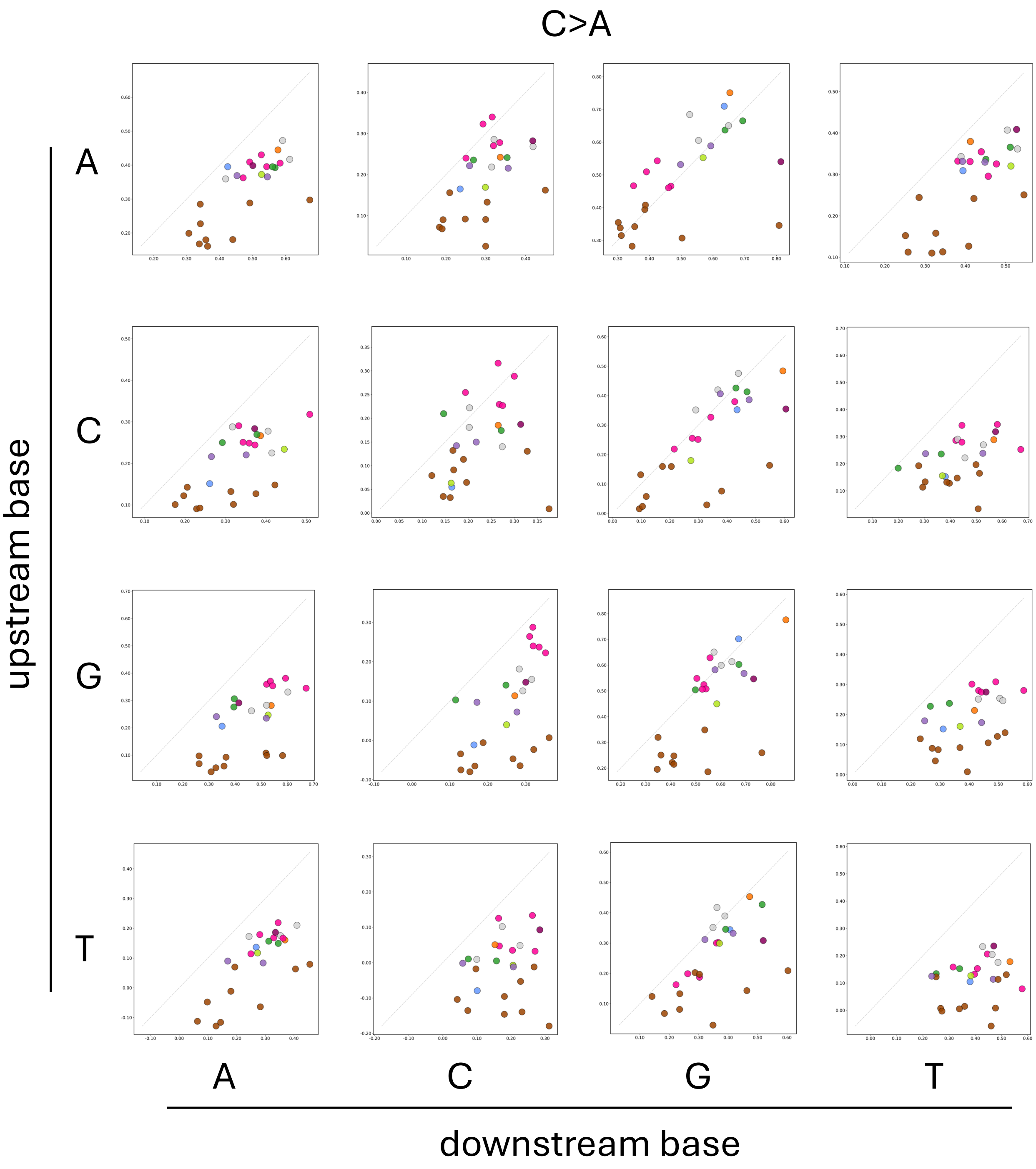


**Fig. S16 | Comparison of reciprocal mutation rate ratios and observed 3-mer frequency ratios for C>A mutations across 25 plant species.**
In each panel, the x-axis shows the ratio of reciprocal mutation rates, and the y-axis shows the corresponding ratio of 3-mer frequencies. Both ratios were $\log_{10}$-transformed. Rows and columns indicate the upstream and downstream bases, respectively. Each point represents one species, colored by group as in Fig. S1.

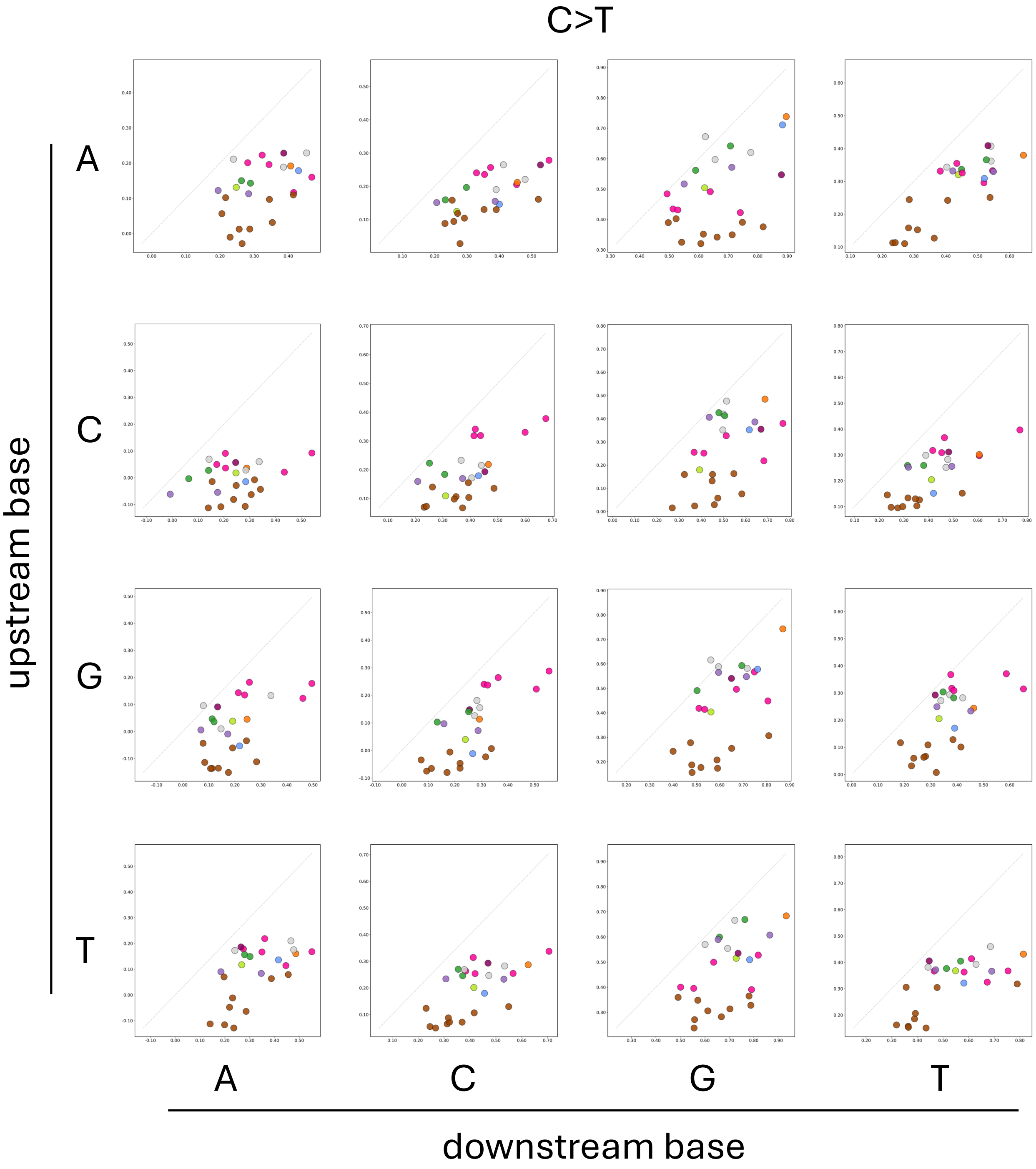


**Fig. S17 | Comparison of reciprocal mutation rate ratios and observed 3-mer frequency ratios for C>T mutations across 25 plant species.**
Same as Fig. S16, but for C>T mutations.

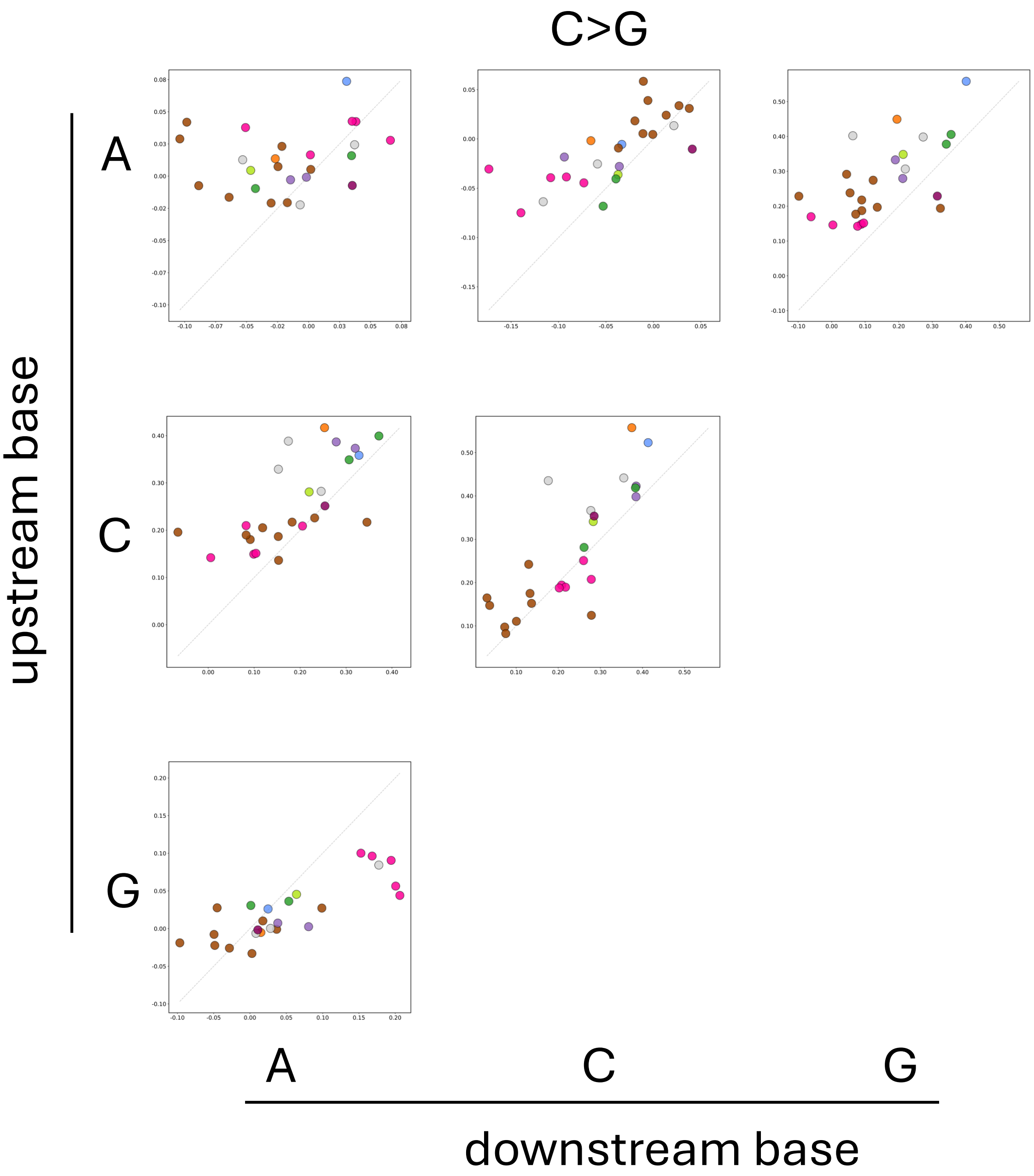


**Fig. S18 | Comparison of reciprocal mutation rate ratios and observed 3-mer frequency ratios for C>G mutations across 25 plant species.**
Same as Fig. S16, but for C>G mutations.

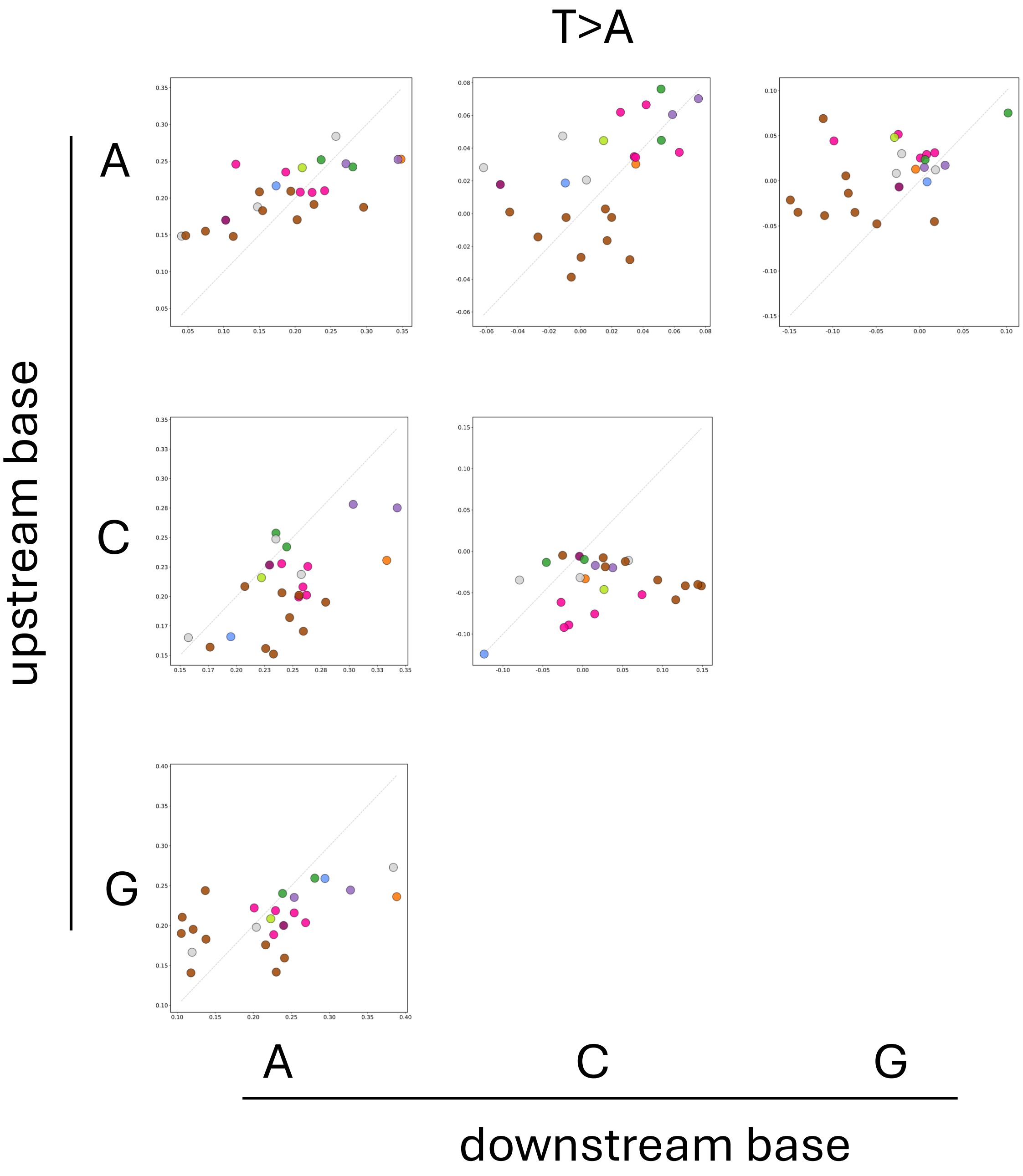


**Fig. S19 | Comparison of reciprocal mutation rate ratios and observed 3-mer frequency ratios for T>A mutations across 25 plant species.**
Same as Fig. S16, but for T>A mutations.

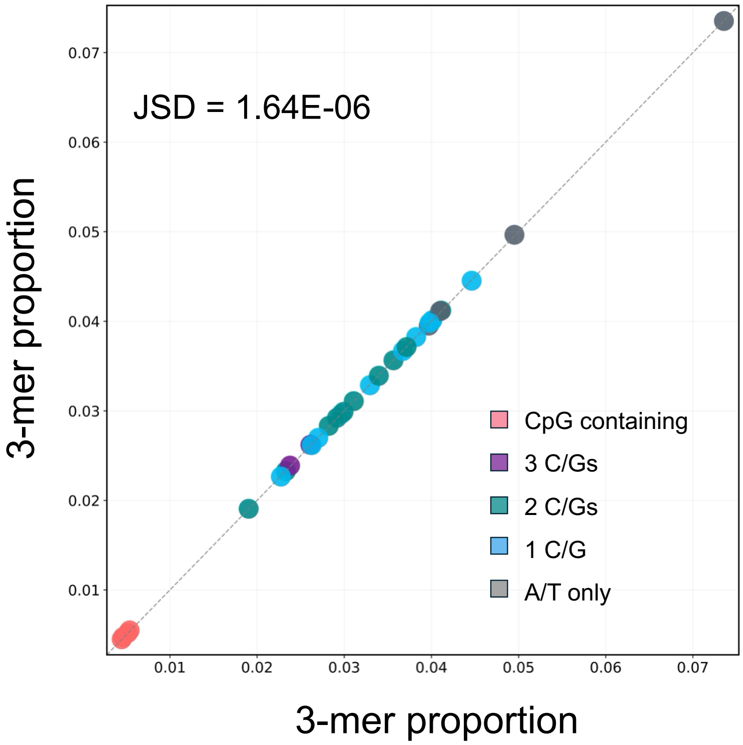


**Fig. S20 | Comparison of noncoding 3-mer frequencies before and after excluding regulatory and conserved regions.**
The scatter plot compares observed 3-mer frequencies in noncoding regions (x-axis) and noncoding regions after excluding candidate regulatory regions (defined as 1kb upstream of TSS and 1kb downstream of TES) and conserved elements (y-axis). See Methods for more details.

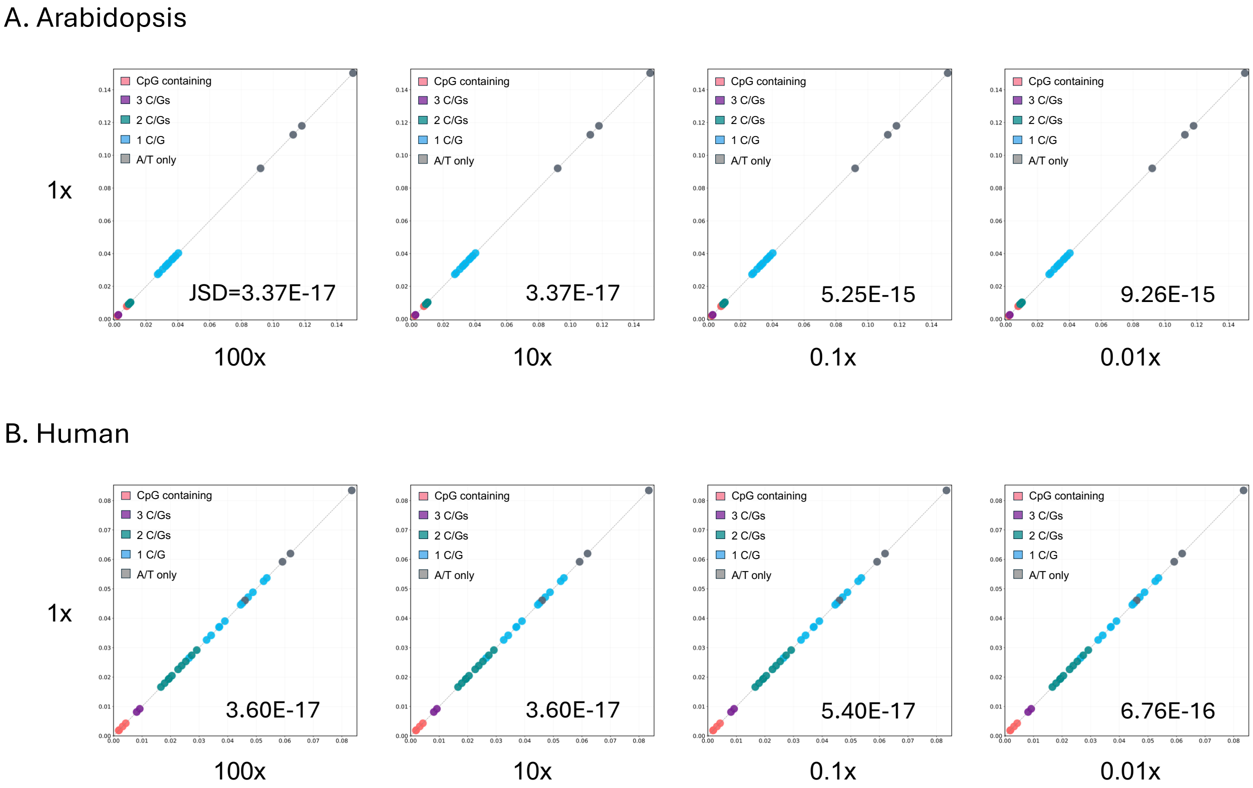


**Fig. S21 | Effect of mutation rate scaling on predicted 3-mer frequencies.**
In *Arabidopsis* (**A**) and human (**B**), scatter plots compare 3-mer frequencies predicted using the 96-channel mutation spectrum from the main analysis (y-axis, see Methods) with those predicted after multiplying all mutation rates by the indicated factor (x-axis).

**Supplementary Tables**

**Table S1. Comparison between predicted and observed genomic composition.**

| Species | JSD (no gBGC) | Predicted GC% (no gBGC) | Observed GC% | Inferred B | JSD (with gBGC) |
|---|---|---|---|---|---|
| Asian_rice | 0.050476 | 27.6% | 42.3% | 0.66 | 0.001757 |
| African_rice | 0.007923 | 35.4% | 41.0% | 0.23 | 0.000998 |
| Wild_rice | 0.076865 | 22.7% | 40.6% | 0.81 | 0.001428 |
| Barley | 0.056665 | 29.4% | 44.4% | 0.72 | 0.002598 |
| Wheat | 0.041140 | 32.9% | 46.0% | 0.57 | 0.002009 |
| Goatgrass | 0.028793 | 35.2% | 46.2% | 0.47 | 0.001187 |
| Foxtail_millet | 0.032809 | 33.3% | 45.1% | 0.52 | 0.000948 |
| Sorghum | 0.024554 | 33.3% | 43.2% | 0.44 | 0.001727 |
| Maize | 0.060592 | 30.7% | 46.7% | 0.69 | 0.003669 |
| Quinoa | 0.019646 | 28.7% | 37.6% | 0.40 | 0.000625 |
| Carrot | 0.003904 | 30.4% | 33.8% | 0.16 | 0.001029 |
| Sticky_monkeyflower | 0.013734 | 26.6% | 33.4% | 0.34 | 0.002013 |
| Alpine_rockcress | 0.046078 | 23.4% | 36.2% | 0.65 | 0.003090 |
| Arabidopsis | 0.036563 | 21.8% | 33.1% | 0.60 | 0.001537 |
| Mustard | 0.007881 | 27.5% | 32.6% | 0.24 | 0.001310 |

| | | | | | |
|---|---|---|---|---|---|
| Cabbage | 0.006224 | 30.0% | 34.6% | 0.21 | 0.001008 |
| Rapeseed | 0.014989 | 27.5% | 34.9% | 0.34 | 0.001133 |
| Cacao | 0.036533 | 21.8% | 33.3% | 0.57 | 0.001437 |
| Cottonwood | 0.020219 | 24.6% | 33.0% | 0.39 | 0.002241 |
| Cassava | 0.033175 | 26.1% | 37.5% | 0.53 | 0.001375 |
| Common_bean | 0.005935 | 30.7% | 35.1% | 0.19 | 0.001312 |
| Soybean | 0.011329 | 28.0% | 34.5% | 0.30 | 0.000963 |
| Hemp | 0.025605 | 24.1% | 34.0% | 0.46 | 0.000785 |
| Apricot | 0.032154 | 25.5% | 36.6% | 0.53 | 0.001056 |
| Peach | 0.003927 | 32.8% | 36.5% | 0.16 | 0.000691 |
| Human | 0.022033 | 34.0% | 41.2% | 0.42 | 0.009390 |
| Chimpanzee | 0.007978 | 36.4% | 40.8% | 0.21 | 0.003420 |
| Mouse | 0.022344 | 33.0% | 42.0% | 0.44 | 0.001891 |
| Porpoise | 0.009689 | 36.5% | 41.6% | 0.24 | 0.003673 |
| Cattle | 0.021988 | 33.2% | 41.9% | 0.43 | 0.003421 |
| Wild_boar | 0.005770 | 38.2% | 42.2% | 0.18 | 0.002045 |
| Dog | 0.040198 | 29.0% | 41.4% | 0.59 | 0.003902 |
| Bear | 0.011956 | 36.6% | 42.4% | 0.29 | 0.004261 |

| | | | | | |
|---|---|---|---|---|---|
| Cat | 0.017295 | 34.1% | 41.8% | 0.37 | 0.003165 |
| Horse | 0.007154 | 38.6% | 42.7% | 0.20 | 0.003169 |

**Table S2. Jensen–Shannon divergence between observed 3-mer proportions and 3-mer proportions predicted based on different models.**

| Species | Mutation only | gBGC | exponential | linear |
|---|---|---|---|---|
| Asian_rice | 0.050476 | 0.001757 | 0.001943 | 0.009953 |
| African_rice | 0.007923 | 0.000998 | 0.001029 | 0.001356 |
| Wild_rice | 0.076865 | 0.001428 | 0.001841 | 0.012869 |
| Barley | 0.056665 | 0.002598 | 0.003101 | 0.010261 |
| Wheat | 0.041140 | 0.002009 | 0.002230 | 0.005093 |
| Goatgrass | 0.028793 | 0.001187 | 0.001410 | 0.002955 |
| Foxtail_millet | 0.032809 | 0.000948 | 0.001093 | 0.003895 |
| Sorghum | 0.024554 | 0.001727 | 0.001865 | 0.002955 |
| Maize | 0.060592 | 0.003669 | 0.003745 | 0.011727 |
| Quinoa | 0.019646 | 0.000625 | 0.000774 | 0.001734 |
| Carrot | 0.003904 | 0.001029 | 0.001080 | 0.001161 |
| Sticky_monkeyflower | 0.013734 | 0.002013 | 0.002101 | 0.002888 |
| Alpine_rockcress | 0.046078 | 0.003090 | 0.003552 | 0.007093 |
| Arabidopsis | 0.036563 | 0.001537 | 0.001813 | 0.003411 |
| Mustard | 0.007881 | 0.001310 | 0.001398 | 0.001425 |
| Cabbage | 0.006224 | 0.001008 | 0.001046 | 0.001151 |
| Rapeseed | 0.014989 | 0.001133 | 0.001250 | 0.001341 |
| Cacao | 0.036533 | 0.001437 | 0.001860 | 0.002893 |
| Cottonwood | 0.020219 | 0.002241 | 0.002462 | 0.002518 |
| Cassava | 0.033175 | 0.001375 | 0.001651 | 0.004353 |
| Common_bean | 0.005935 | 0.001312 | 0.001350 | 0.001257 |
| Soybean | 0.011329 | 0.000963 | 0.001053 | 0.001021 |
| Hemp | 0.025605 | 0.000785 | 0.000855 | 0.001852 |
| Apricot | 0.032154 | 0.001056 | 0.001318 | 0.002633 |
| Peach | 0.003927 | 0.000691 | 0.000724 | 0.000703 |
| Human | 0.022033 | 0.009390 | 0.009688 | 0.012674 |
| Chimpanzee | 0.007978 | 0.003420 | 0.003496 | 0.003872 |
| Mouse | 0.022344 | 0.001891 | 0.002228 | 0.003362 |
| Porpoise | 0.009689 | 0.003673 | 0.003670 | 0.004222 |
| Cattle | 0.021988 | 0.003421 | 0.003793 | 0.005213 |
| Wild_boar | 0.005770 | 0.002045 | 0.002116 | 0.002262 |
| Dog | 0.040198 | 0.003902 | 0.004311 | 0.009184 |
| Bear | 0.011956 | 0.004261 | 0.004295 | 0.005381 |
| Cat | 0.017295 | 0.003165 | 0.003384 | 0.004898 |
| Horse | 0.007154 | 0.003169 | 0.003236 | 0.003665 |

**Table S3. Jensen–Shannon divergence between observed 3-mer proportions and those predicted using 6-channel, 7-channel, 9-channel and 96-channel mutation spectra.**

| Species | 6-channel | 7-channel | 9-channel | 96-channel |
|---|---|---|---|---|
| Asian_rice | 0.004915 | 0.006104 | 0.006279 | 0.001757 |
| African_rice | 0.004789 | 0.005231 | 0.005331 | 0.000998 |
| Wild_rice | 0.004832 | 0.003652 | 0.005919 | 0.001428 |
| Barley | 0.006479 | 0.006718 | 0.006578 | 0.002598 |
| Wheat | 0.005444 | 0.009229 | 0.009021 | 0.002009 |
| Goatgrass | 0.005045 | 0.007933 | 0.007913 | 0.001187 |
| Foxtail_millet | 0.005213 | 0.005711 | 0.005615 | 0.000948 |
| Sorghum | 0.005475 | 0.003662 | 0.003634 | 0.001727 |
| Maize | 0.005772 | 0.005001 | 0.005026 | 0.003669 |
| Quinoa | 0.007840 | 0.005940 | 0.005935 | 0.000625 |
| Carrot | 0.005841 | 0.003467 | 0.003349 | 0.001029 |
| Sticky_monkeyflower | 0.006060 | 0.004486 | 0.004570 | 0.002013 |
| Alpine_rockcress | 0.006443 | 0.005892 | 0.005902 | 0.003090 |
| Arabidopsis | 0.006762 | 0.005626 | 0.005522 | 0.001537 |
| Mustard | 0.005551 | 0.004864 | 0.004850 | 0.001310 |
| Cabbage | 0.005446 | 0.005240 | 0.005225 | 0.001008 |
| Rapeseed | 0.005573 | 0.005327 | 0.005356 | 0.001133 |
| Cacao | 0.011102 | 0.007016 | 0.006536 | 0.001437 |
| Cottonwood | 0.010936 | 0.008509 | 0.008353 | 0.002241 |
| Cassava | 0.011103 | 0.007199 | 0.006371 | 0.001375 |
| Common_bean | 0.009283 | 0.006848 | 0.006635 | 0.001312 |
| Soybean | 0.010239 | 0.006773 | 0.006457 | 0.000963 |
| Hemp | 0.006988 | 0.004905 | 0.004607 | 0.000785 |
| Apricot | 0.010388 | 0.006837 | 0.006821 | 0.001056 |
| Peach | 0.009711 | 0.006649 | 0.006529 | 0.000691 |
| Human | 0.031236 | 0.009461 | 0.008724 | 0.009390 |
| Chimpanzee | 0.024372 | 0.008586 | 0.007698 | 0.003420 |
| Mouse | 0.029798 | 0.008448 | 0.005595 | 0.001891 |
| Porpoise | 0.022864 | 0.009800 | 0.008431 | 0.003673 |
| Cattle | 0.031674 | 0.010104 | 0.008452 | 0.003421 |
| Wild_boar | 0.024983 | 0.009593 | 0.007622 | 0.002045 |
| Dog | 0.033634 | 0.010334 | 0.008184 | 0.003902 |
| Bear | 0.023505 | 0.011132 | 0.009851 | 0.004261 |
| Cat | 0.023270 | 0.011299 | 0.010090 | 0.003165 |
| Horse | 0.023429 | 0.010378 | 0.009512 | 0.003169 |

**Table S4: Pearson's r values for within-species correlations between mutation-rate ratios and 3-mer proportion ratios.**

| Species | no gBGC | with gBGC | observed |
|---|---|---|---|
| Asian_rice | 0.906 | 0.631 | 0.619 |
| African_rice | 0.951 | 0.888 | 0.871 |
| Wild_rice | 0.974 | 0.691 | 0.824 |
| Barley | 0.798 | 0.538 | 0.431 |
| Wheat | 0.840 | 0.605 | 0.557 |
| Goatgrass | 0.882 | 0.666 | 0.648 |
| Foxtail_millet | 0.804 | 0.588 | 0.579 |
| Sorghum | 0.903 | 0.767 | 0.611 |
| Maize | 0.956 | 0.579 | 0.155 |
| Quinoa | 0.958 | 0.865 | 0.823 |
| Carrot | 0.956 | 0.951 | 0.914 |
| Sticky_monkeyflower | 0.985 | 0.957 | 0.917 |
| Alpine_rockcress | 0.862 | 0.809 | 0.668 |
| Arabidopsis | 0.857 | 0.836 | 0.747 |
| Mustard | 0.934 | 0.931 | 0.898 |
| Cabbage | 0.965 | 0.962 | 0.945 |
| Rapeseed | 0.963 | 0.933 | 0.900 |
| Cacao | 0.926 | 0.858 | 0.821 |
| Cottonwood | 0.929 | 0.891 | 0.727 |
| Cassava | 0.931 | 0.869 | 0.825 |
| Common_bean | 0.989 | 0.965 | 0.952 |
| Soybean | 0.992 | 0.953 | 0.944 |
| Hemp | 0.958 | 0.899 | 0.816 |
| Apricot | 0.947 | 0.854 | 0.836 |
| Peach | 0.987 | 0.975 | 0.970 |
| Human | 0.794 | 0.741 | 0.634 |
| Chimpanzee | 0.946 | 0.937 | 0.929 |
| Mouse | 0.975 | 0.953 | 0.954 |
| Porpoise | 0.944 | 0.933 | 0.924 |
| Cattle | 0.942 | 0.913 | 0.902 |
| Wild_boar | 0.990 | 0.986 | 0.987 |
| Dog | 0.934 | 0.879 | 0.868 |
| Bear | 0.931 | 0.910 | 0.883 |
| Cat | 0.953 | 0.929 | 0.925 |
| Horse | 0.956 | 0.944 | 0.931 |

The "no gBGC" column indicates the predicted composition without gBGC; the "with gBGC" column indicates the predicted composition with gBGC; and the "observed" column indicates the observed composition in noncoding regions.

**Table S5: Pearson's r values for cross-species correlations between mutation-rate ratios and 3-mer proportion ratios.**

| Mutation type | Pearson's *r* | *P* | adjusted *P* |
|---|---|---|---|
| A[C>A]A | 0.728 | 0.079 | 1.000 |
| A[C>A]C | 0.500 | 0.229 | 1.000 |
| A[C>A]G | 0.593 | 0.238 | 1.000 |
| A[C>A]T | 0.702 | 0.274 | 1.000 |
| C[C>A]A | 0.646 | 0.104 | 1.000 |
| C[C>A]C | 0.299 | 0.969 | 1.000 |
| C[C>A]G | 0.704 | 0.343 | 1.000 |
| C[C>A]T | 0.417 | 0.933 | 1.000 |
| G[C>A]A | 0.613 | 0.067 | 1.000 |
| G[C>A]C | 0.577 | 0.099 | 1.000 |
| G[C>A]G | 0.662 | 0.109 | 1.000 |
| G[C>A]T | 0.455 | 0.322 | 1.000 |
| T[C>A]A | 0.719 | 0.115 | 1.000 |
| T[C>A]C | 0.132 | 0.377 | 1.000 |
| T[C>A]G | 0.564 | 0.126 | 1.000 |
| T[C>A]T | 0.220 | 0.890 | 1.000 |
| A[C>G]A | 0.223 | 0.364 | 1.000 |
| A[C>G]C | 0.715 | 0.564 | 1.000 |
| A[C>G]G | 0.612 | 0.309 | 1.000 |
| G[C>G]A | 0.818 | 0.270 | 1.000 |
| G[C>G]G | 0.808 | 0.009 | 0.389 |
| T[C>G]G | 0.681 | 0.026 | 1.000 |
| A[C>T]A | 0.442 | 0.690 | 1.000 |
| A[C>T]C | 0.630 | 0.075 | 1.000 |
| A[C>T]G | 0.459 | 0.055 | 1.000 |
| A[C>T]T | 0.818 | 0.173 | 1.000 |
| C[C>T]A | 0.279 | 0.276 | 1.000 |
| C[C>T]C | 0.648 | 0.095 | 1.000 |
| C[C>T]G | 0.501 | 0.855 | 1.000 |
| C[C>T]T | 0.694 | 0.057 | 1.000 |
| G[C>T]A | 0.501 | 0.372 | 1.000 |
| G[C>T]C | 0.703 | 0.071 | 1.000 |
| G[C>T]G | 0.571 | 0.028 | 1.000 |
| G[C>T]T | 0.613 | 0.773 | 1.000 |
| T[C>T]A | 0.560 | 0.396 | 1.000 |
| T[C>T]C | 0.627 | 0.191 | 1.000 |
| T[C>T]G | 0.512 | 0.109 | 1.000 |

| | | | |
|---|---|---|---|
| T[C>T]T | 0.652 | 0.248 | 1.000 |
| A[T>A]A | 0.711 | 0.073 | 1.000 |
| A[T>A]C | 0.492 | 0.201 | 1.000 |
| A[T>A]G | 0.390 | 0.761 | 1.000 |
| C[T>A]A | 0.603 | 0.078 | 1.000 |
| C[T>A]C | 0.239 | 0.266 | 1.000 |
| G[T>A]A | 0.570 | 0.267 | 1.000 |

All 44 pairs used in Table S4 are listed here. In Fig. 4E, C[C>G]A and T[C>G]G, as well as C[C>G]C and G[C>G]G, are reverse-complement pairs and therefore have identical Pearson's *r*. The "*P*" column reports the *P* values from phylogenetic regression, whereas the "adjusted *P*" column reports the corresponding *P* values after Bonferroni correction for multiple testing.

**Supplementary datasets**

**Dataset S1.** Genome assembly information and repeat-masking statistics for the species included in the comparative genomic analyses. The dataset includes species classification, genome version and source, assembly level, genome size before and after masking, and the proportion of the genome masked.

**Dataset S2.** Proportions of trinucleotides in intergenic regions for each species analyzed in this study, with each trinucleotide combined with its reverse-complementary counterpart.

**Dataset S3.** Relative mutation spectra for each species analyzed in this study. Mutation rates were normalized within each species so that the sum across all mutation types equaled 1.

**Dataset S4.** Proportions of trinucleotides in noncoding regions for each species analyzed in this study, with each trinucleotide combined with its reverse-complementary counterpart.

**Dataset S5.** Genome assembly source information for the additional 13 plant species included in the phylogenetic tree construction (See Methods).
